# The Arabidopsis RS2Z32 and RS2Z33 proteins are dynamic splicing factors whose RNA recognition motif (RRM) domain contributes to protein-protein and protein-RNA interactions

**DOI:** 10.1101/2024.09.04.610270

**Authors:** Steven Fanara, Marie Schloesser, Méline Gérard, Simona De Franco, Marylène Vandevenne, Marc Hanikenne, Patrick Motte

## Abstract

The Arabidopsis splicing factors arginine/serine-rich zinc knuckle-containing proteins 32 and 33 (RS2Z32 and RS2Z33) are plant-specific members of the SR family whose molecular functions received little attention. Here, we characterized both RS2Z32 and RS2Z33 by examining their expression profile at different stages of development and their spatial cellular distribution, as well as the contribution of their domains in the establishment of protein-protein interactions and RNA binding specificity. We report that the *RS2Z32* and *RS2Z33* promoters are ubiquitously active during vegetative and reproductive growth, and that both RS2Z splicing factors localize in the nucleus (except the nucleolus). We show that the C-terminal arginine/serine-rich (RS) domain, but not the serine/proline-rich (SP) extension, is a determinant of nuclear localization, which likely requires phosphoresidues putatively phosphorylated by kinases of the SRPK family. We demonstrate that their RNA recognition motif (RRM) domain specifically binds pyrimidine-rich RNA motifs via three residues (Y14, Y46, F48), and is also involved in protein-protein interactions with at least three SR proteins, namely SR45, SCL30, and SR34. Finally, we show that mutations in RNA-binding domains (i.e. RRM and zinc knuckles, ZnKs) affect the nucleocytoplasmic dynamics of both RS2Z proteins. Our findings provide molecular evidence for the involvement of plant-specific SR splicing factors into the regulation of the splicing process.

**Highlight:** Specific domains of the *Arabidopsis* RS2Z splicing factors contribute to their nuclear localization, nucleocytoplasmic dynamics, and ability to contact protein partners and specific pyrimidine-rich RNA motifs.

## Introduction

The precursor messenger RNA (pre-mRNA) splicing process requires the precise recognition of splice sites at exon-intron boundaries. It is achieved by the spliceosome, consisting of small nuclear ribonucleoproteins and a large number of non-snRNP-associated proteins (Meyer *et al.,* 2015). If constitutive splicing produces a single transcript from the pre-mRNA by excluding all introns, alternative splicing (AS) generates several mRNAs from a single precursor sequence through different events (Marquez *et al.,* 2015). These events require the recognition of non-canonical (e.g., weak) splice sites by the splicing machinery and may drive transcriptomic, hence proteomic diversity (Manuel *et al.,* 2023), or lead to nonsense-mediated mRNA decay (NMD) due to premature termination codons (PTCs) occurring in aberrant alternative transcripts (Chen and Manley 2009; Palusa and Reddy 2010; Kalyna *et al.,* 2012). Native and alternative protein isoforms usually diverge in their primary sequence or domain organization, which could alter the interaction network established with protein, DNA or RNA, as well as the protein function, stability, or intracellular localization (English *et al.,* 2010; Kalyna *et al.,* 2012; Drechsel *et al.,* 2013; Remy *et al.,* 2013; Kashkan *et al.,* 2022).

Serine/arginine-rich (SR) splicing factors are phylogenetically highly conserved in multicellular eukaryotes (Califice *et al.,* 2012) whose members share a modular structure defined by one or two N-terminal RNA recognition motif(s) [RRM(s)] and a C-terminal domain rich in arginine-serine dipeptide repeats (RS) (Barta *et al.,* 2010; Manley and Krainer 2010; Califice *et al.,* 2012). The RRM domain(s) support(s) the RNA binding affinity and specificity (Maris *et al.,* 2005; Daubner *et al.,* 2013; Fanara *et al.,* 2024). The RS domain undergoes extensive reversible phosphorylation (Aubol *et al.,* 2016; Stankovic *et al.,* 2016; Aubol *et al.,* 2017; Fanara *et al.,* 2024). The phosphorylation landscape of the RS domain modulates the ability of SR proteins to contact protein partners, as well as the nucleocytoplasmic dynamics and the subcellular localization of SR proteins, notably by contributing to their phospho-dependent transport into the nucleus via an SR-specific transportin protein (TRN-SR, TNPO3) (Kataoka *et al.,* 1999; Lai *et al.,* 2000; Lai *et al.,* 2001; Xu *et al.,* 2011). A phylogenetic analysis of RRM-containing proteins highlighted the existence of nineteen SR protein-encoding genes in the *Arabidopsis thaliana* (Arabidopsis) genome (Califice *et al.,* 2012). These proteins are subdivided into seven groups, including four plant-specific subfamilies (RS, SCL, RS2Z and SR45) each defined by a unique multidomain organization (Barta *et al.,* 2010; Califice *et al.,* 2012).

The arginine/serine-rich zinc knuckle-containing splicing factors (RS2Z) bear two CCHC-type zinc knuckles (hereafter referred as ZnK1 and ZnK2, and ZnKs if discussed simultaneously) located between a unique RRM and the RS domain, as well as a C-terminal serine/proline-rich (SP) domain of unknown function **(Supplementary Figure S1)** (Barta *et al.,* 2010; Califice *et al.,* 2012). The overexpression of *RS2Z33* was shown to lead to increased cell expansion, a modified polarization of cell elongation and division, and an altered auxin signalling (Kalyna *et al.,* 2003). The involvement of RS2Z33 in cell patterning and root meristem function was recently further corroborated, depicting a role for RS2Z33 in primary and lateral root development (Thompson *et al.,* 2023). Both *RS2Z* genes undergo alternative splicing events leading to NMD features (e.g., PTC upstream of a splice junction) (Kalyna *et al.,* 2012; Thompson *et al.,* 2023). Post-translational modifications of native RS2Z isoforms were previously reported, including ubiquitylation (Ub-conjugates) (Kim *et al.,* 2013), methylation of an arginine residue (R80) in the RS2Z32, but not RS2Z33, protein (Liang *et al.,* 2019) and phosphorylation of both proteins by members of the SRPKII family (SRPK3, SRPK4 and SRPK5) (Wang *et al.,* 2023). The inventory of protein partners was well detailed for RS2Z33 and regroup many splicing factors, including SR34, RSZ21, RSZ22, SC35, SCL28, SCL30, and SCL33 (Lopato *et al.,* 2002), as well as MOS14 (Xu *et al.,* 2011), but not RZ-1B or RZ-1C (Wu *et al.,* 2016). The Cyclin-Dependent Kinase G1 (CDKG1) (Huang *et al.,* 2013) and the cyclophilin AtCyp59 (Gullerova *et al.,* 2006) are also partners of RS2Z33. Recently, SR45 was identified as a novel interactor of both RS2Z32 and RS2Z33 (Fanara *et al.,* 2024).

In Arabidopsis, RS2Z32 and RS2Z33 proteins remain poorly characterized (e.g., ability to shuttle from the nucleus to the cytoplasm, or nature of the RNA motif bound). In this study, we aimed to characterize both proteins through the investigation of their spatio-temporal expression pattern, their subcellular distribution and nucleocytoplasmic dynamics, and their protein-protein interaction network. We further determined critical amino acid residues supporting these functions. We also identified the RNA motif bound by the RRM domain of each RS2Z proteins, observing slight discrepancies in the motifs specifically recognized by each protein, and demonstrated how three amino acids (Y14, Y46, F48) define their RNA binding specificity. Finally, we showed that phosphorylatable serines are necessary and sufficient for establishing the RS2Z protein-protein interaction network. Our findings provide new insights into the molecular characterization of plant-specific splicing factors.

## Materials and methods

### Plant material and growth conditions

*Arabidopsis thaliana* plants (Col-0 ecotype) were stably transformed by floral dipping, and at least three independent T3 homozygous lines were analyzed for each construct. Seeds were surface-sterilized, then sown on 1/2 Murashige and Skoog (MS) medium (Duchefa Biochimie) supplemented with sucrose (1% w/v, Duchefa Biochimie) and Select Agar (0.8% w/v, Sigma-Aldrich), and stratified in the dark at 4°C for 48h. Petri dishes were then incubated in a climate-controlled growth chamber (21°C) under an 8-h light (100 μmol m^-2^ sec^-1^)/16-h dark regime. 3- week-old seedlings were transferred in hydroponic trays (Araponics, Liège, Belgium) and cultivated for 7 weeks in control Hoagland medium (Talke *et al.,* 2006; Hanikenne *et al.,* 2008) (until silique development) under a 16-h light (100 μmol m^-2^ sec^-1^)/8-h dark regime in a climate-controlled growth chamber (21°C). Expression profiling was conducted on 10-week-old plants hydroponically grown. The nutrient medium was renewed with fresh solution once a week.

*Nicotiana tabacum* (cv Petit Havana) transient transformations by *Agrobacterium* infiltration were performed as described (Rausin *et al.,* 2010; Stankovic *et al.,* 2016).

### Vector constructions

All binary vector constructions and primers used in this study are listed and detailed in **Supplementary Tables S1-S8**. The Q5 DNA polymerase (NEB) was used on Arabidopsis cDNA libraries or genomic DNA to amplify all gene and promoter sequences, respectively. All constructs were verified by sequencing. For plant transient and stable transformations, all final plasmids were electroporated into the *Agrobacterium tumefaciens* strain GV3101 (pMP90) and used for agroinfiltration in tobacco leaves or Arabidopsis floral dipping, respectively.

For directed yeast two-hybrid assays (dY2H), cDNAs coding for the potential interactors were cloned into pGBKT7/pGBKT7(+) and pGADT7/pGADT7(+) vectors (Clontech) using different restriction sites (**Supplementary Table S1** for primer lists) (Fanara *et al.,* 2024).

The promoter regions of *RS2Z32* and *RS2Z33* [701 bp and 1110 bp upstream of the ATG, respectively] were amplified by PCR from genomic DNA. The promoter amplicons were ligated at the *Sbf*I/*Kpn*I sites of the pMDC32 vector (Curtis and Grossniklaus 2003) to create the pMDC32:pRS2Z vectors. The *RS2Z32* or *RS2Z33* open reading frame (852 bp and 870 bp, from the ATG to the last codon before the stop codon, respectively) was then cloned at *Asc*I/*Pac*I sites to obtain pMDC32:pRS2Z:RS2Z intermediate constructs. Finally, the *EGFP* coding sequence was cloned at the *Pac*I site to create the pMDC32:pRS2Z:RS2Z:EGFP vectors. To construct the pMDC32:pRS2Z:EGFP vectors, the 2X35S promoter of the pMDC32 vector (Curtis and Grossniklaus 2003) was replaced by the *RS2Z32* or *RS2Z33* promoter and EGFP was cloned downstream of the promoter at *Asc*I and *EcoR*I restriction sites **(Supplementary Table S2)**.

The *RS2Z32mutRRM* and *RS2Z33mutRRM* versions were obtained by PCR-based site-directed mutagenesis as described (Rausin *et al.,* 2010; Stankovic *et al.,* 2016) **(Supplementary Table S3)**. The other mutated *RS2Z* versions were assembled by PCR from partially overlapping amplicons, which corresponded to the coding sequence of each domain of RS2Z32 or RS2Z33. They were amplified using either the native *RS2Z32* or *RS2Z33* genes, the corresponding *RS2ZmutRRM* gene, or the RS or SP mutated domains (named RA and AP) synthesized by GenScript (see **Supplementary Tables S4 and S5**). All assembled mutant versions **(Supplementary Figure S1)** were then introduced into yeast vectors for Y2H (at *Eco*RI/*Bam*HI) or pMDC32 fused to EGFP for protein localization (at *Asc*I/*Pac*I sites), respectively.

The BiFC constructs were obtained as described (Stankovic *et al.,* 2016; Fanara *et al.,* 2024) **(Supplementary Table S6)**.

For SELEX experiments, genes encoding the RRM or RRM+ZnKs of RS2Z32 or RS2Z33 were cloned at *Bam*HI/*Eco*RI sites of the pGEX6P-1 as described (De Franco *et al.,* 2019) **(Supplementary Table S7)**.

### Yeast two-hybrid analyses

The vectors and yeast (*Saccharomyces cerevisiae*) strains provided in the Matchmaker Gold Yeast Two-Hybrid System (Clontech) were used to perform directed interaction analyses (dY2H) through co-transformation of the Y2HGold strain. Prior to (directed) Y2H experiments, toxicity and autoactivation of the wild-type *RS2Z32* or *RS2Z33* and mutant variants were tested simultaneously on a medium -Trp/X-a-gal [for pGBKT7] and -Leu/X-a-gal [for pGADT7] using 100 ng of plasmid. Yeast growth was not affected by any of the constructs, none of which led to autoactivation of the reporter gene (*MEL1*) **(Supplementary Figure S2)**. Dimeric interactions were then tested on selective medium -Trp/-Leu/-His/X-a-gal/Aureobasidin A.

For Y2H screening, the Y2HGold strain was transformed with the bait vector (pGBKT7- RS2Z32 or pGBKT7-RS2Z33) and then mated with strain Y187 containing an Arabidopsis cDNA library (Mate and Plate Library-Universal Arabidopsis, Clontech) or a custom cDNA library made from entire mature leaf material (Make Your Own “Mate & Plate” Library System, Clontech) (Fanara *et al.,* 2024).

### Confocal Microscopy

Live cell imaging was performed on Leica TCS SP2 and SP5 inverted confocal laser microscopes (Leica Microsystems). A water-immersion objective was used to collect images at 512x512 pixel resolution. EGFP was visualized using the 488-nm line of the argon laser, and the emission light was dispersed and recorded at 500 to 560 nm (Fanara *et al.,* 2024). For BiFC experiments, YFP was excited at 514 nm using the argon laser and the emission light was recorded at 525 to 600 nm, and the voltage of the EYFP PMT was set up at 700-900 V for isolated nuclei observations, and at 800-900V for maximum intensity projections of 30 z-stack images (Fanara *et al.,* 2024). The FLIP-shuttling experiments were carried out and analyzed as previously described (Tillemans *et al.,* 2006; Rausin *et al.,* 2010; Stankovic *et al.,* 2016) **(Supplementary Table S8)**.

### Analysis of the mRNA levels

Total RNAs were extracted from 100 mg of plant tissues using NucleoSpin RNA Plant kit (Macherey Nagel). 1 µg of total RNAs was used to synthesize cDNAs using the RevertAid H Minus First Strand cDNA Synthesis Kit (ThermoFisher Scientific) with oligo(dT) primers. Quantitative PCR reactions were performed in 384-well plates using a QuantStudio5 system (Applied Biosystems) and Takyon Low Rox SYBR MasterMix dTTP Blue (Eurogentec). Samples were obtained from three independent biological experiments. A total of three technical replicates were performed for each combination of cDNA and primer pair **(Supplementary Table S9)**. Gene expression was normalized relative to *At1g58050* as described (Pfaffl 2001). *At1g58050* expression was the most stable gene among all tested references (*EF1α* and *UBQ10*) (Czechowski *et al.,* 2005; Spielmann *et al.,* 2020).

### Production of recombinant proteins

The recombinant protein production was performed using the expression vector pGEX6P- 1 in *E. coli* BL21(DE3) cells as described (De Franco *et al.,* 2019). Protein productions (native and mutated RRM of RS2Z32 or RS2Z33, or native RRM+ZnKs of RS2Z32 or RS2Z33) were carried out overnight at 18°C upon induction with 0.5 mM IPTG and supplemented with 100 µM ZnCl_2_. Cells were lysed from sonication in 50 mM Tris-HCl pH 8, 1 M NaCl, 2 mM MgCl_2_, 100 µM ZnCl_2_, 1 mM dithiothreitol (DTT) and 1 mM PMSF. GST fusion proteins were purified on Glutathione-Sepharose^®^ 4B beads (GE Healthcare) and eluted using a solution of 50 mM Tris-HCl pH 8, 100 mM NaCl, 100 µM ZnCl_2_, 1 mM dithiothreitol (DTT) and 35 mM reduced glutathione.

### SELEX experiments

The Systematic Evolution of Ligands by EXponential enrichment (SELEX) experiment was performed using a randomized 25-nt DNA template library as previously reported (De Franco *et al.,* 2019; Fanara *et al.,* 2024). *In vitro* transcription reaction (T7 RiboMAX™ Express Large Scale RNA Production System, Promega) was performed using 500 ng-1 µg of the purified template DNA library. Mini Quick Spin RNA Columns (Roche) were used to remove unincorporated nucleotides. RNAs were purified with phenol/chloroform and then ethanol-precipitated. The RNA pellet was resuspended in RNAse-free water, and the final RNA concentration was quantified by measuring the absorbance at 260/280 nm. Binding reactions were carried out in 200 µL of SELEX buffer (20 mM MOPS pH 7.0, 50 mM KCl, 5 mM MgCl_2_, 100 µM ZnCl_2_, 1 mM DTT, 0.1% Triton-X-100, and 0.1 mM PMSF) and included 80 pmol of GST-fused protein immobilized on GSH beads (GE Healthcare), 1–8 µg of heparin sulfate, and 0.8–4 nmol of RNAs. Samples were gently agitated at 4 °C for 60 min. Unbound RNA was removed by five consecutive bead washings in 500 µL of SELEX buffer. HCl-25 mM glycine (pH 2.2) was used to dissociate RNA-protein complexes at 25 °C for 5 min, and the dissociated RNAs were subsequently purified using mini Quick Spin RNA Columns (Roche). They were then ethanol-precipitated, and reverse-transcribed (SuperScript® III Reverse Transcriptase, Invitrogen). The resulting cDNAs were PCR amplified by 10 cycles for SELEX rounds 1 to 3, or 15 cycles for last SELEX round. The amplified cDNAs were then purified (PCR clean up kit, Thermo Fisher Scientific) and used as a template to synthesize a new RNA library for the following SELEX rounds. The selection stringency was intensified by increasing the RNA/protein ratios and the heparin concentration in every SELEX round. The final PCR products (round 4) were subcloned into the pJET vector (CloneJET PCR Cloning Kit, Thermo

Fisher Scientific) and sequenced **(Supplementary Tables S10-15)**. The obtained sequences were analyzed using the MEME tool (version 5.5.1) (Bailey *et al.,* 2009). Logos were redesigned using WebLogo software (Crooks *et al.,* 2004).

### Statistical analysis

All data evaluation and statistics were performed using GraphPad Prism 8 (GraphPad Software v8.0). The replication level of each experiment is described in the figure legends. For statistical analysis of FLIP-shuttling assays, the fluorescence intensities at three given time points [10s, 65s (50% of the time scale) or 129s] of each FLIP-shuttling experiment were processed as follow: a statistical analysis of normality was performed using the D’Agostino and Pearson test. To calculate the significance of the differences between fluorescence intensities, an unpaired t test (parametric data) was performed. The Wilcoxon signed-rank test was used when at least one of the series of data failed the normality test (Stankovic *et al.,* 2016). The differences observed were considered to be statistically significant if *p*-values were at least < 0.05, as indicated by asterisks (*P < 0.05, **P < 0.01, ***P < 0.001, ****P < 0.0001). n.s., not significant.

### Accession numbers and information

All gene sequences are available through The Arabidopsis Information Resource (TAIR, http://www.arabidopsis.org/), with the following accession numbers: Arabidopsis *ACINUS* (AT4G39680), *AFC1* (AT3G53570), *AFC2* (AT4G24740), *AFC3* (AT4G32660), *ALY4* (AT5G37720), *Cyp59* (AT1G53720), *Cyp65* (AT5G67530), *Cyp71* (AT3G44600), *CypRS64* (AT3G63400), *CypRS92* (AT4G32420), *eIF4A3* (AT3G19760), *GRP7* (AT2G21660), *GRP8* (AT4G39260), *MAGO* (AT1G02140) *MOS12* (AT2G26430), *MOS14* (AT5G62600), *PININ* (AT1G15200), *PRP38* (AT2G40650), *RS2Z232* (AT3G53500), *RS2Z33* (AT2G37340), *RS31* (AT3G61860), *RS31a* (AT2G46610), *RS40* (AT4G25500), *RS41* (AT5G52040), *RSZ21* (AT1G23860), *RSZ22* (AT4G31580), *RSZ22a* (AT2G24590), *RZ-1B* (AT1G60650), *RZ-1C* (AT5G04280), *SAP18* (AT2G45640), *SC35* (AT5G64200), *SCL28* (AT5G18810), *SCL30* (AT3G55460), *SCL30a* (AT3G13570), *SCL33* (AT1G55310), *SR30* (AT1G09140), *SR34* (AT1G02840), *SR34a* (AT3G49430), *SR34b* (AT4G02430), *SR45.1* (AT1G16610.1), *SR45.2* (AT1G16610.2), *SRPK1* (AT4G35500), *SRPK2* (AT2G17530), *SRPK3a* (AT5G22840), *SRPK3b/SRPK5* (AT3G44850), *SRPK4* (AT3G53030), and *Y14* (AT1G51510).

## Results

### RS2Z genes encode nuclear proteins and are expressed at all developmental stages

Quantitative RT-PCR profiling showed that *RS2Z32* and *RS2Z33* genes were expressed in all vegetative (roots, leaves, stems and seedlings) and reproductive (inflorescences and siliques) organs examined during plant development. However, *RS2Z32* was significantly more expressed compared to *RS2Z33* in all organs but roots, with a significant greater expression for both genes in siliques and young seedlings **(Supplementary Figure S3)**. This corroborates and largely deepens the previous quantification available for these genes (Palusa *et al.,* 2007; Cruz *et al.,* 2014). The expression profile of both *RS2Z* genes and the subcellular and tissular distribution of the two splicing factors were also further determined using promoter:reporter lines [*pRS2Z32:EGFP* **(Supplementary Figure S4A-M)** or *pRS2Z33:EGFP* **(Supplementary Figure S4a-m)**] and GFP translational fusions [*pRS2Z32:RS2Z32-EGFP* **(Figure 1A-Q)** or *pRS2Z33:RS2Z33-EGFP* **(Figure 1a-q)**], respectively. Translational fusion lines displayed a nuclear localization of both RS2Z members in the seed coat **(Figure 1A/a)** and in embryos freshly dissected from seeds **(Figure 1B/b)**. During vegetative growth, the hypocotyl and cotyledons displayed fluorescence **(Figure 1C/c-D/d)**, which was also detected in roots and root hairs **(Figure 1E/e-F/f)**, as well as leaves, trichomes and stomata **(Figure 1G/g-I/i)**. Immature floral buds were dissected to reveal intense fluorescence signals in the receptacle, the abscission zone, sepals, petals, stamens and the pistil **(Figure 1J/j)**. During flowering, mature styles and stigmas emitted fluorescence signals **(Figure 1K/k)**, as well as stamens (anthers, filaments and pollen grains) **(Figure 1L/l-N/n)**. GFP fluorescence also resolved in ovules and funiculi **(Figure 1O/o)**, and fully developed petals and sepals (including veins) **(Figure 1P/p-Q/q)**. The observations in reporter lines were in agreement with those of the translational fusion lines, and also largely complete previous results depicting the activity of the promoter of *RS2Z33* in the embryo, leaves, roots, ovules and pollen grains (Kalyna *et al.,* 2003) **(Supplementary Figure S4A/a-M/m)**.

**Figure 1.**
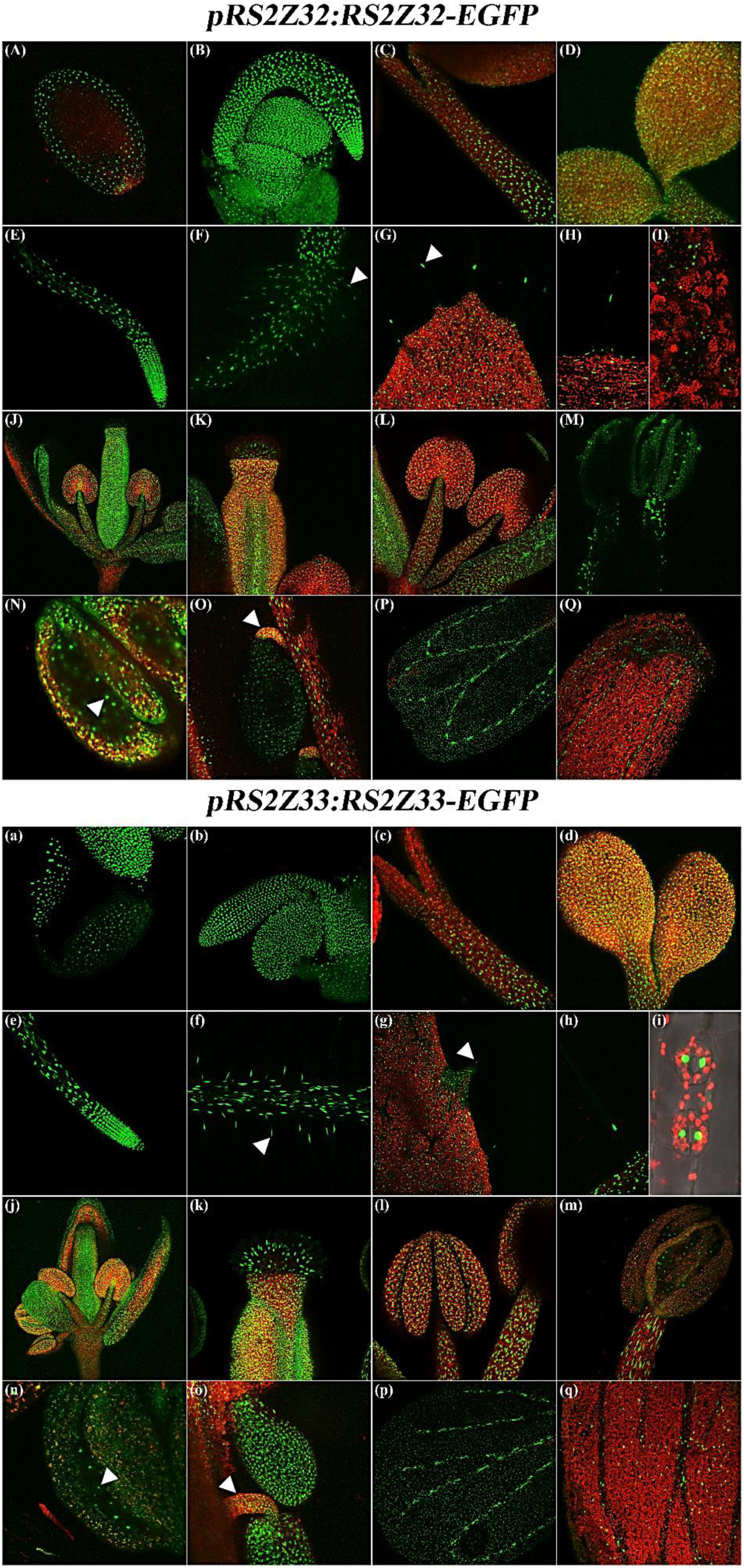
Localization of RS2Z proteins in all vegetative and reproductive organs. Translational fusions are expressed in seed coat **(A/a)**, embryo **(B/b)**, hypocotyl **(C/c)** and cotyledons **(D/d),** two-week-old root epidermis **(E/e)** including root hairs **(F/f)**, leaf epidermis **(G/g-I/i)** including trichomes **(G/g-H/h)** and stomata **(I/i)**, immature floral bud (including sepals, petals, stamens and pistil) **(J/j)**, mature gynoecium (valves, stigma and style) **(K/k)**, mature androecium (anther and filament) **(L/l)**, pollen grains [**(M/m-N,n)**, arrow], ovules **(O/o)** and funiculi [**(O/o)**, arrow)], petals (including veins) **(P/p)** and sepals (including veins) **(Q/q)**. Red signals represent chlorophyll autofluorescence. At least three independent T3 homozygous lines were generated and analyzed, depicting similar fluorescence profiles.

### RS2Z localization depends on native serine residues within the RS domain

The subcellular localization of the two RS2Z proteins, as well as mutant variants for each of their domains, was next examined. Twelve mutant variants were engineered **(Supplementary Figure S1)**. First, the RS2ZmutRNP2 and RS2ZmutRNP1 variants were obtained by substituting to alanines conserved residues mediating the RNA specificity [Y14 (RNP2), Y46 (RNP1) and F48 (RNP1)] of the canonical RRM (Cléry *et al.,* 2008), and combining these substitutions allowed the generation of the RS2ZmutRRM variants. Second, all cysteine residues from each CCHC-type zinc knuckle were substituted into alanine residues to generate the RS2ZmutZnK1 and RS2ZmutZnK2 variants, whose mutations were combined to obtain the RS2ZmutZnKs variants. Mutations within the RRM and both ZnKs were merged to produce the RS2ZmutRRM+ZnKs variants. Third, all Ser residues were substituted to Ala within the RS (17 S-to-A substitutions) and/or SP domains (15 and 16 S-to-A substitutions for RS2Z32 and RS2Z33, respectively), generating three variants (mutRS, mutSP, mutRS+SP), respectively. Finally, RRM and RS or SP mutations were combined, producing two more mutant variants (mutRRM+RS and mutRRM+SP) **(Supplementary Figure S1)**.

The intracellular localization of the native and variant RS2Z proteins was determined after transient expression in tobacco leaf cells, confirming the localization of RS2Z32 and RS2Z33 within the nucleoplasm and speckles-like structures but not the nucleolus (Lorković and Barta 2004; Tillemans *et al.,* 2005). No distinguishable changes appeared in the protein localization of RS2ZmutRRM (or lower order mutant variants; mutRNP1 and mutRNP2), RS2ZmutZnKs (or lower order mutant variants; mutZnK1 and mutZnK2), RS2ZmutRRM+ZnKs, RS2ZmutSP and RS2ZmutRRM+SP mutant variants compared to the native protein **(Figure 2 and Supplementary Figure S5)**. In contrast, RS2ZmutRS, RS2ZmutRRM+RS and RS2ZmutRS+SP mutant variants partially accumulated in the cytoplasm in addition to a nuclear and nucleolar localization. However, the RS2ZmutRRM+RS displayed a fainter cytoplasmic signal compared with the two other mutant variants bearing the mutated RS **(Figure 2)**. Altogether, the presence of native serine residues within the RS domain appears necessary to ensure a strict nucleoplasmic localization of both RS2Z32 and RS2Z33.

**Figure 2.**
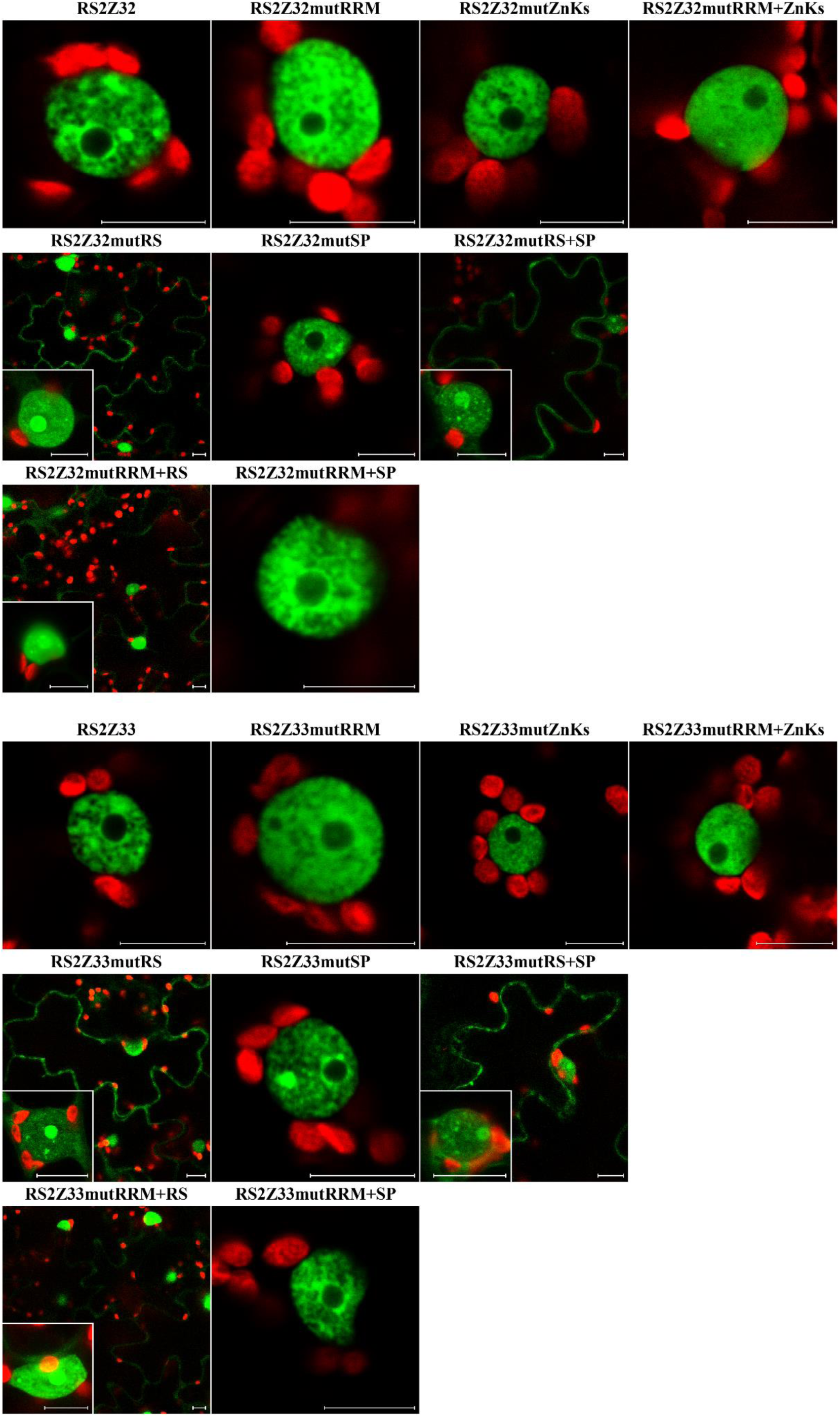
Structural determinants modulating the localization of RS2Z members. Subcellular fluorescence distribution in transient expression assays in tobacco leaf cells upon N-terminal EGFP-tagging of the native RS2Z32 or RS2Z33 and their respective mutant variants. Scale bars = 10 µm. Red signals represent chlorophyll autofluorescence. Whenever cytoplasmic and nuclear signals were detected, insets depict the detailed intranuclear localization. At least three independent transient events were generated and analyzed, depicting similar fluorescence profiles.

### RS2Z dynamics depends on RNP2 motif and of ZnK2 motif integrity and is partly controlled by XPO1

We previously determined that RSZ22, SR34 and SR45 were shuttling splicing factors whose nuclear export might be partly controlled by the XPO1-dependent export pathway (Tillemans *et al.,* 2006; Rausin *et al.,* 2010; Stankovic *et al.,* 2016). For the SR34 protein (Stankovic *et al.,* 2016), we further showed that the canonical RRM (RRM1) contributes to its nucleocytoplasmic dynamics. On the contrary, the canonical RRM or ZnK of RSZ22 are dispensable for its nucleocytoplasmic shuttling (Rausin *et al.,* 2010). In this context, we aimed to determine whether (i) the RS2Z proteins displayed a shuttling behavior, (ii) whether it may be dependent on the XPO1 export pathway, and (iii) whether the RRM and/or the ZnKs contributed to the functional shuttling of these factors. To this avail, we specifically characterized the dynamics of RS2Z32 and RS2Z33 using FLIP-shuttling assays, focusing on the seven mutants harboring mutations within the RNP motifs of the RRM and/or within the CCHC-type zinc knuckle(s) **(Supplementary Figure S1)**. The native RS2Z proteins were highly dynamic factors, as indicated by the rapid loss of fluorescence upon photobleaching **(Figure 3A/a)**. Treatment with leptomycin B (LMB), a XPO1-specific inhibitor, led to a statistically significant lower shuttling kinetics, suggesting a XPO1-dependent nuclear export **(Figure 3A/a)**. While mutations in the RNP1 or ZnK1 did not drastically affect the shuttling activity of the RS2Z proteins in absence of LMB treatment **(Figure 3B/b and D/d, Supplementary Table S8)**, alterations of the RNP2 significantly impacted the dynamics of these splicing factors, indicated by a lower shuttling kinetics **(Figure 3C/c, Supplementary Table S8)**. Substitutions in the ZnK2 mildly but significantly impaired the shuttling activity of RS2Z33, but no significant modification of the RS2Z32mutZnK2 dynamics was observed compared to the native RS2Z32 protein **(Supplementary Table S8)**. Combining either RNP1 and RNP2 or ZnK1 and ZnK2 mutations (i.e. mutRRM and mutZnKs, respectively) did not aggravate the impairment of the shuttling dynamics of these mutant variants **(Supplementary Figure S6A/a-B/b)**. Interestingly, while the RS2ZmutRNP1 variant behaved in the same fashion as the native RS2Z protein upon LMB treatment **(Figure 3B/b, Supplementary Table S8)**, the RS2ZmutRNP2 variant was statistically significantly less sensitive to LMB treatment, displaying a rapid rate of fluorescence loss similarly to what was observed without LMB treatment **(Figure 3C/c, Supplementary Table S8)**. The dynamic behavior of the ZnK1 mutant variant was less impacted by LMB treatment **(Figure 3D/d)**, displaying a rather fast decrease of fluorescence, but the ZnK2 mutant variant was drastically affected by the inhibitor in an identical fashion compared to the native protein submitted to LMB **(Figure 3E/e Supplementary Table S8)**. The ZnKs mutant variant displayed an intermediate behavior regarding the effect of LMB on its dynamics **(Supplementary Figure S6B/b)**. The RS2ZmutRRM+ZnKs variant remained dynamic, and its shuttling activity was mildly impacted by LMB treatment **(Supplementary Figure S6C/c, Supplementary Table S8)**.

**Figure 3.**
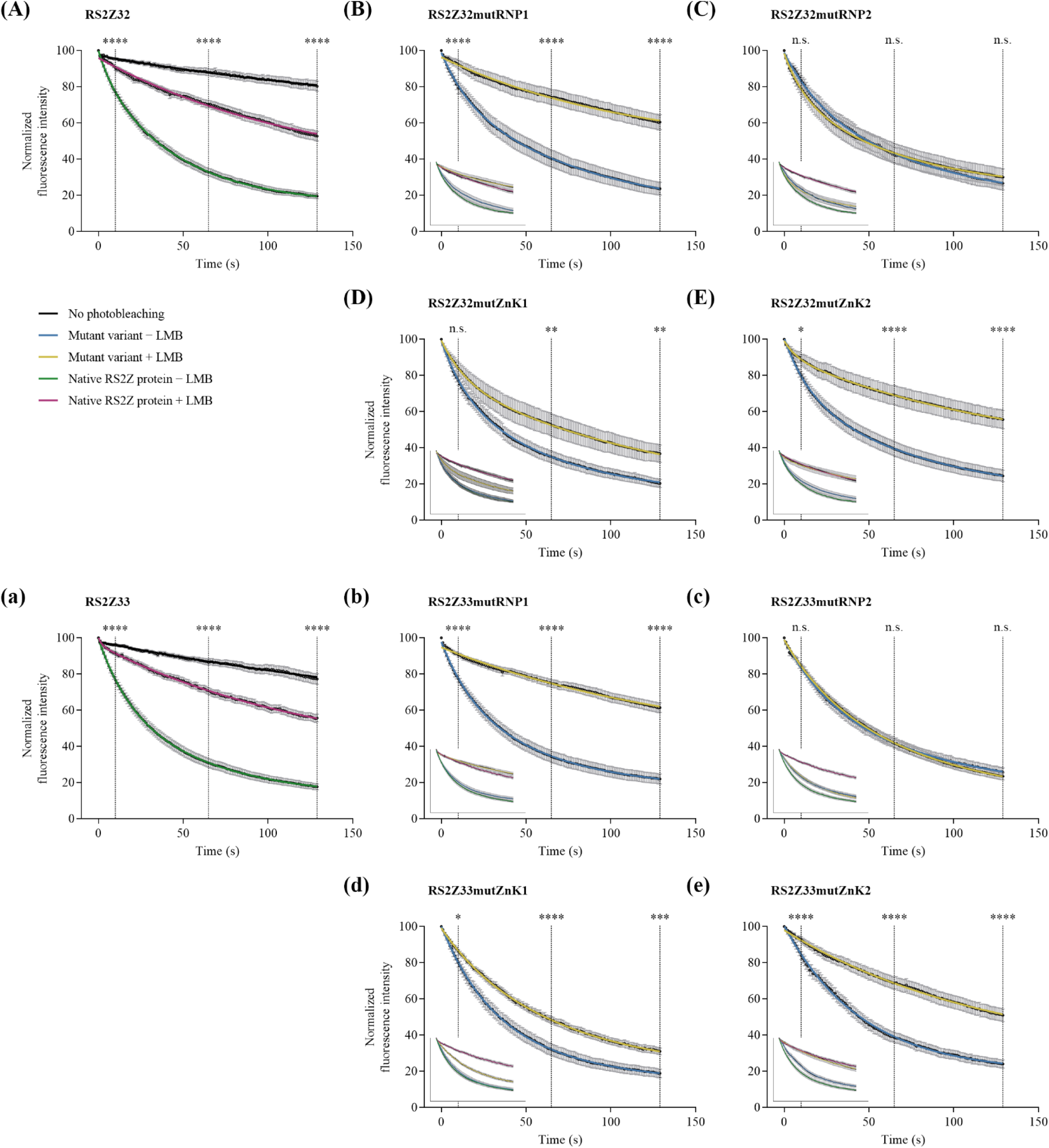
Nucleocytoplasmic shuttling of native RS2Z proteins and mutant variants. FLIP-shuttling was assessed without (−LMB) or with LMB (+LMB) treatment in tobacco epidermal leaf cells. One hundred percent fluorescence indicates prebleach fluorescence intensity. As a control of the dynamic nature of native RS2Z proteins, cells were repeatedly scanned under no photobleaching conditions and fluorescence was monitored. Insets show the overlay of wild-type and mutant curves. Values are means ± SEM for at least 10 nuclei. A significant inhibitory effect of LMB is indicated by asterisks (*P < 0.05, **P < 0.01, ***P < 0.001, ****P < 0.0001). n.s., not significant. Three given time points [10s, 65s (50% of the time scale) or 129s] were statistically analyzed.

### The RRM of RS2Z proteins contributes to contacting splicing factors

In a previous work (Lopato *et al.,* 2002), a yeast two-hybrid (Y2H) screen was performed to identify RS2Z33 protein interactors. It used a truncated version of the protein (RS2Z33ΔSP) due to the yeast toxicity of the full-length protein. In this screen, RS2Z33ΔSP was found to interact with SR34, RSZ21, RSZ22, SC35, SCL28, SCL30, and SCL33 (Lopato *et al.,* 2002). As the native untruncated RS2Z32 protein was recovered during a recent Y2H screen using SR45 as a bait (Fanara *et al.,* 2024), we used here full-length versions of the two RS2Z proteins in new Y2H analyses using two Arabidopsis cDNA libraries (a commercially available and a custom-made) in yeast two-hybrid (Y2H) assays. This led to the identification of previously unidentified partners of RS2Z32 (SCL30 and CypRS64), as well as known partners of RS2Z33 (SCL30, SR45.1 and SR45.2), confirming the SCL30-RS2Z33 association previously reported (Lopato *et al.,* 2002). Directed yeast two-hybrid assays (dY2H) allowed the confirmation of these interactions **(Figure 4A, Supplementary Figure S7 and Supplementary Figure S8)**.

**Figure 4.**
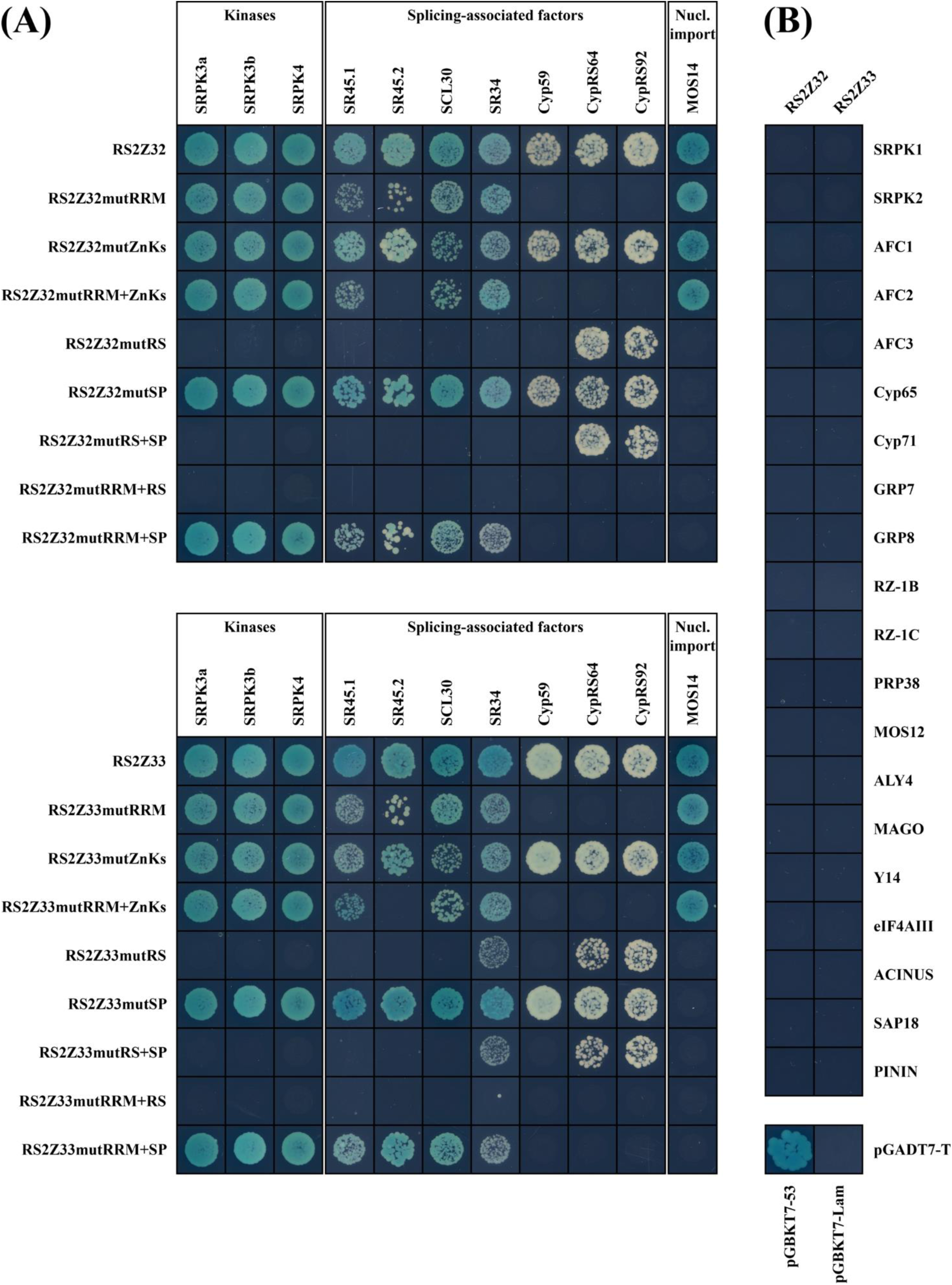
RS2Z32 and RS2Z33 associates with kinases, splicing-associated factors, and the nuclear importer. **(A)** The baits tested correspond to native and mutant variants. **(B)** Absence of interaction between RS2Z32 or RS2Z33 with kinases (SRPK1, SRPK2, and AFC1-3), nuclear cylophilins (Cyp65 and Cyp71), hnRNP-like proteins (GRP7, GRP8, RZ-1B, and RZ-1C), PRP38, MOS12, ALY/REF ortholog ALY4, EJC core components (MAGO, Y14, eIF4AIII), or the peripheral EJC components (ACINUS, PININ, and SAP18). From the initial culture, dilutions to an OD600 of 0.25 were spotted on -Trp/-Leu/-His/X-α-Gal/AurA agar plates. Positive interactions were confirmed by growth and blue staining as seen with the positive control (pGADT7-T x pGBKT7-53). An abolished interaction is characterized at most by a shadow of dead cells (similar to the negative control pGADT7-T×pGBKT7-Lam). Serial dilutions (OD 0.25, 0.025 and 0.0025) are presented in **Supplementary Fig. S7**.

The dY2H set-up was further used to examine associations between all Arabidopsis SR proteins and the native RS2Z proteins. This highlighted strong contacts between RS2Z proteins and both SR45 isoforms (Zhang and Mount 2009). Contrary to RS2Z interactions with RSZ21, SR34a, SCL28 or SCL30a, which appeared to be relatively weak, RS2Z also strongly associated to SR34, SCL30 and SCL33 **(Supplementary Figure S8A-B)**. Upon testing all mutant variants engineered in this study, the RS2Z-SR45 isoforms contacts were weakened or disrupted upon RRM and RS mutations, respectively **(Figure 4A)**, however, mutant harboring both altered RRM and ZnKs domains were not able to contact the SR45.2 isoform contrary to what was observed with the SR45.1 isoform **(Figure 4A)**. Moreover, increasing serial dilutions did not allow the growth of yeast cells, attesting of the weaken interaction occurring between RS2ZmutRRM variants and the SR45.2 isoform **(Supplementary Figure S7)**. We further showed that the sole RS2Z RRM (RS2ZΔZnKs+RS+SP) sustained a weakened interaction with SR45 isoforms **(Supplementary Figure S8C)**, and that the RS2Z proteins were predominantly contacted by the RS1 domain of SR45.1 **(Supplementary Figure S9)**. Moreover, SR45.1 RS1 mutations impacted more strongly the connection to RS2Z32 compared to RS2Z33 **(Supplementary Figure S9)**. A complete abolition of these contact was observed upon mutations applied to both RS domains of SR45.1 **(Supplementary Figures S9 and S10)**. Similarly, the RS2Z-SCL30 interactions were also weakened or abolished upon mutations applied to the RRM (or even to ZnKs) and to the RS domain, respectively. Mutations of the RS domain of RS2Z32 and RS2Z33, respectively, abolished and weakened the association to SR34. The RS2Z33-SR34 association was fully disrupted when combining RS and RRM mutations **(Figure 4A)**.

Kinases potentially involved in the phosphorylation of serine residues of the RS and SP domains were also sought using dY2H. While none of the 3 *cdc2*-like kinases (CLK) kinases (AFC1-3) (Bender and Fink 1994) interacted with the RS2Z proteins **(Figure 4B)**, 3 out of 5 SRPK kinases (SRPK3a, SRPK3b [also named SRPK5] and SRPK4 (Wang *et al.,* 2023)) were found to strongly interact with RS2Z32 and RS2Z33, which confirmed a recent report (Wang *et al.,* 2023) **(Figure 4A and Supplementary Figure S7)**. Mutations applied to the RS domain abolished the association to SRPK3a, SRPK3b and SRPK4 **(Figure 4A)**. Those results suggested that the RS domain, but not the SP, is involved in interactions with kinases of the SRPK family, arguing that phosphorylation of serine residues in the RS domain may take place.

As nuclear cyclophilins are involved in pre-mRNA maturation and are known to interact with SR proteins (Lorković and Barta 2004; Gullerova *et al.,* 2006; Barbosa Dos Santos and Park 2019; Jo *et al.,* 2022), we tested the associations of RS2Z proteins to Cyp59, Cyp65, Cyp71, CypRS64 and CypRS92 **(Figure 4A-B)**. Cyp59, CypRS64 and CypRS92 were found to interact with the RS2Z members, however, the mutRRM variants were unable to sustain these contacts. Mutations affecting the RS domain were also responsible for the complete disruption of the RS2Z-Cyp59 association, hence only the native proteins and the mutant variants harboring altered ZnKs or SP domains were able to contact Cyp59 **(Figure 4A)**. The interaction between Cyp59 and RS2Z32 appeared to be weaker than the one involving RS2Z33, as depicted by the lack of resilience of yeast cells to grow upon serial dilutions (also noticeable at the starting OD of 0.25). **(Supplementary Figure S7)**.

It was shown that RS2Z33 interacts with MOS14, a transportin-SR (TRN-SR) thought to drive the nuclear import of SR proteins (Xu *et al.,* 2011). Here, we found that both RS2Z32 and RS2Z33 were able to interact with MOS14, which required an intact RS domain, as well as an intact SP domain **(Figure 4A)**.

Finally, no interaction was observed between hnRNP-like proteins (GRP7, GRP8, RZ-1B, and RZ-1C), PRP38, MOS12, ALY/REF ortholog ALY4, EJC core components (MAGO, Y14, eIF4AIII), or the peripheral EJC components (ACINUS, SAP18, and PININ) and the full-length versions of RS2Z proteins **(Figure 4B)**.

Our observations depict the involvement of serine residues in contacting kinases and splicing factors, which also requires in some extent the presence of a native canonical RRM (containing unaltered RNP1 and RNP2 motifs).

### RS2Z proteins strongly interacts with mRNA processing factors in planta

As the RS2Z proteins interact with few SR proteins that are known to interact between themselves (e.g., SCL30-SR34 (Lorković *et al.,* 2008), SR34-SR45 (Stankovic *et al.,* 2016), and SR45-SCL30 (Fanara *et al.,* 2024)) and that are all contacted by CypRS64 (Lorković *et al.,* 2004; Fanara *et al.,* 2024; this study), we used Yellow Fluorescent Protein (YFP)-based bimolecular fluorescence complementation (BiFC) to confirm *in planta* interactions observed in yeast cells. RS2Z proteins fused to the C-terminal fragment of YFP (^C^YFP) and interactors fused to the N-terminal fragment of YFP (^N^YFP) were transiently co-expressed in tobacco leaf cells. This allowed to determine that RS2Z proteins interacted with SR45.1, SCL30, SR34 and CypRS64 within the nucleoplasm and inside speckles-like granules in tobacco leaf cells **(Figure 5)**. The interaction with SR45 was dependent on its RS domains **(Supplementary Figure S10)**.

**Figure 5.**
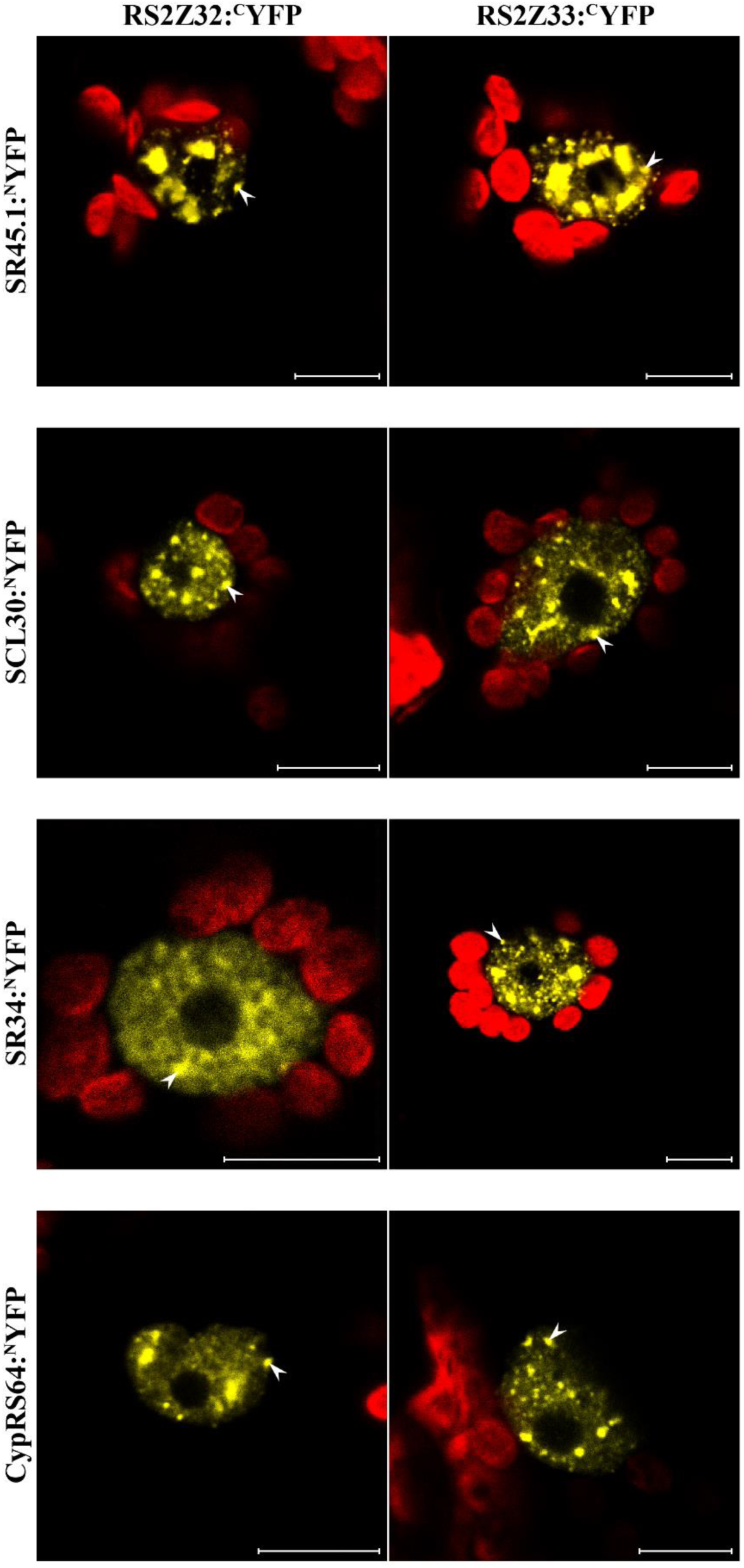
RS2Z proteins interact with mRNA processing factors. Bimolecular fluorescence complementation (BiFC) in transient expression assay in tobacco leaf cells through co-expression of RS2Z32 or RS2Z33 fused to the C-terminal half of YFP (RS2Z32:^C^YFP or RS2Z33:^C^YFP) and either **(A)** SR45.1., SCL30, SR34 and CypRS64 fused to the N-terminal half of YFP (SR45.1, SCL30, SR34, or CypRS64:^N^YFP). For all observations, representative images of fluorescence reveal the interactions of RS2Z32 or RS2Z33 with the respective proteins in the nucleus, and in speckle-like structures (arrowheads). At least three independent transient events were generated and analyzed, depicting similar fluorescence profiles. Scale bars = 10 µm. Red signals represent chlorophyll autofluorescence.

### RNA binding specificity of RNA-contacting domains of the RS2Z proteins

We then determined the RNA sequences bound by the RS2Z factors. Recombinant RRM domains of RS2Z32 and RS2Z33 fused to a glutathione-S-transferase were used in a Systematic Evolution of Ligands by EXponential enrichment (SELEX) experiment. Among the 10 independent clones sequenced after SELEX and used during the MEME analysis, a common central motif, 5’-G[G/C]UA-3’, flanked by variable regions was found to be recognized by the two RS2Z proteins **(Figure 6A-B)**. While both RRM recognized a U/C-rich stretch at the 3’-end of their respective RNA motif, the 5’-end was highly variable, with a striking displacement of an invariable ‘C’ nucleotide, from the position 4 in the RNA motif for RS2Z32 to the position 3 for RS2Z33. We therefore studied the occurrence of ‘CC’ dinucleotides in the sequences used by MEME to identify each motif, which showed that 60% and 71% of the sequences contained the motif 5’-CCGGUA-3’ for RS2Z32 and RS2Z33, respectively. Moreover, most of the sequences used by MEME contained the central motif 5’-GGUA-3’ instead of the 5’-GCUA-3’ (80% and 71% of the cases for RS2Z32 and RS2Z33, respectively) **(Supplementary Tables S10-11)**.

**Figure 6.**
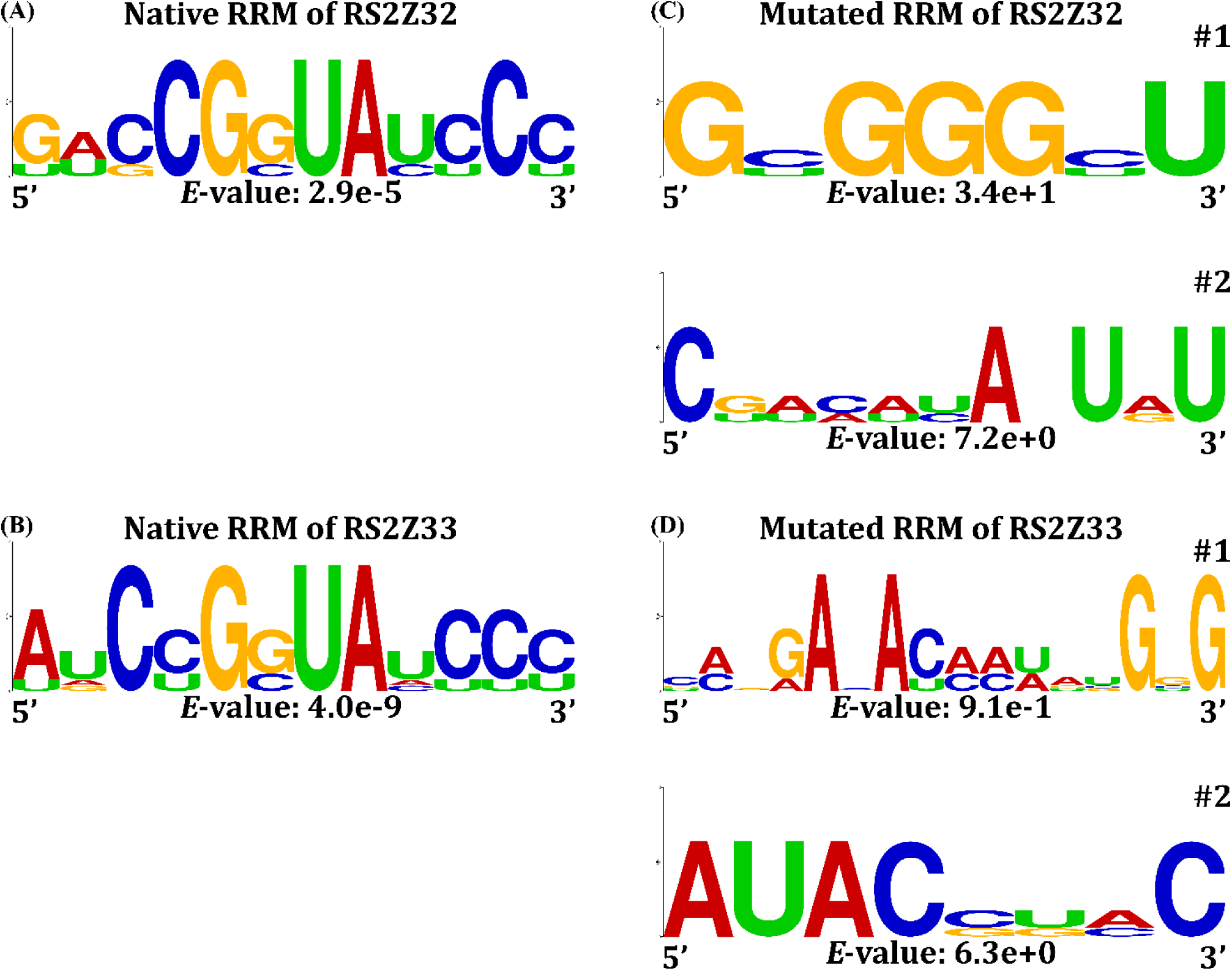
The RRM of RS2Z proteins binds to a pyrimidine-rich RNA motif. RNA motifs were identified through 4 rounds of SELEX selection using either the native RRM of RS2Z32 **(A)** and RS2Z33 **(B)** or the mutated versions of the RRM of RS2Z32 **(C)** and RS2Z33 **(D)**. Two non-significant RNA motifs (#1-2) were identified using the mutated versions. The statistical significance (*E*-value) is indicated at the bottom of each consensus. Logos were redesigned using WebLogo (Crooks *et al.,* 2004). All RNA motifs were discovered using the MEME tool (version 5.5.1) with 10 sequences.

Substitutions to alanine of three conserved residues (Y14, and Y46 and F48) thought to mediate RNA recognition and specificity disrupted the recovery of a significant motif by SELEX **(Figure 6C-D)**. Indeed, two different sets of ten sequences obtained with the mutated RS2Z32 or RS2Z33 RRM were analyzed **(Supplementary Tables S12-13)** using the MEME tool, and no enriched motif was discovered **(Figure 6C-D).** Interestingly, among the sequences selected through binding to the native RRM of RS2Z32 and RS2Z33, purine and pyrimidine contents were even (with a 2.5-3% enrichment in purine bases) **(Supplementary Tables S10-11)**. In contrast, upon Y14A-Y46A-F48A substitutions, purine bases were enriched by ∼14% to ∼22% on average in sequences contacted by RS2Z32 and RS2Z33 **(Supplementary Tables S12-13)**. This also confirmed the importance of these residues in the selectivity of sequences able to be connected by the native RRM.

Although Zn knuckles were shown to contribute to the RNA binding specificity of many splicing factors (De Franco *et al.,* 2019), including SRSF7 (Cavaloc *et al.,* 1999; Königs *et al.,* 2020), SELEX experiments performed using recombinant proteins including both RRM and ZnKs did not uncover any significant RNA motif **(Supplementary Figure S11 and Supplementary Tables S14-15)**. Altogether, our results demonstrate that the amino acids Y14, Y46 and F48 of the RRM govern the RNA binding specificity of each RS2Z protein for a pyrimidine-stretch-containing motif.

## Discussion

Among the many steps orchestrating pre-mRNA processing, splicing is a crucial maturing event in the regulation of gene expression. Many *trans*-acting protein factors, including SR splicing factors, are involved in splice-site selection through the recognition of and binding to specific *cis*-elements (Chen and Manley 2009). Mammalian SR proteins were extensively studied and are known to regulate many aspects of gene expression (Huang and Steitz 2001; Long and Cáceres 2009). The functional characterization of mammalian SR proteins involved the study of their constitutive domains, such as the ZnK of the human ortholog of the RSZ (one RRM and one ZnK) subfamily (SRSF7) which has been found to interact with multiple RNA sequences (Cavaloc *et al.,* 1999). While the study of their plant RSZ counterparts is less detailed, the functional relationship between RRM and ZnK was previously explored in the context of the nucleocytoplasmic shuttling activity and subcellular localization of RSZ22 (Rausin *et al.,* 2010). Contrary to the plant SR45 protein, whose mammalian ortholog (RNPS1) is well characterized, which favoured assumptions about the SR45 protein-protein network and molecular functions (Fanara *et al.,* 2024), RS2Z32 and RS2Z33 belong to a plant-specific subfamily with a characteristic presence of a canonical RRM domain, directly followed by two CCHC-type zinc knuckles (ZnKs), and Arg/Ser-rich (RS) and Ser/Pro-rich (SP) domains (Barta *et al.,* 2010). To this date, the characterization of this subfamily mainly focused on one of the RS2Z at a time (Lopato *et al.,* 2002; Kalyna *et al.,* 2003) but was never conducted in a comparative fashion.

Using mutant variants, we evaluated how each of the RRM, ZnKs, RS and SP domains (e.g., their constituent amino acid residues) modulate the molecular functions played by the RS2Z proteins. To achieve this goal, three point mutations were introduced in conserved residues of the RRM domain (Y14A, Y46A, F48A), both CCHC-type ZnKs were altered by Cys-to-Ala substitutions, and all serine residues of the RS or SP domains were substituted by alanines. The impact on the global structure of RS2Z proteins and functionality of all mutant variants constructed in this study was evaluated by assessing several parameters defining a typical SR splicing factor, i.e. (i) their intracellular localization; (ii) their nucleocytoplasmic dynamics, showing that shuttling kinetics were differently affected depending on the mutations ; (iii) the protein–protein interactions, highlighting that different mutations abolished the association to different partners; and (iv) the protein–RNA interactions, proving the existence of contacts between the mutated RRM and RNA molecules despite an alteration of specificity (Yang *et al.,* 2011; Pabis *et al.,* 2019; Ni *et al.,* 2023; Fanara *et al.,* 2024). While we showed changes in the molecular functions of the engineered mutant variants compared to the native RS2Z proteins, these changes were likely not linked to the impairment of structure of the domains mutated as the SR proteins are known to be poorly structured outside of the RNA recognition motif (RRM) (Haynes and Iakoucheva 2006), therefore an unstructured region would not be affected upon alanine substitutions. Collectively, these observations dismiss the possibility of a complete non-functionality of the engineered mutant variants.

### Alteration of the RS domain of RS2Z proteins leads to their nucleolar sequestration and disrupt their ability to contact splicing factors

We observed that the RS domain of both RS2Z proteins often reinforces the contact between their RRM and splicing factors. Indeed, the substitution of all serine within the RS domain (RS2ZmutRS) by alanine residues can either severely diminish or even abolish the contact between splicing partners (e.g., SR45 isoforms, SCL30 and SR34) and these RS2Z mutant variants. Interestingly, S-to-A substitutions within the RS domain also affect the subcellular localization of the RS2Z members, which leads to nucleolar sequestration and to a partial cytoplasmic accumulation of the RS2ZmutRS variant suggesting a role for the RS domain in the control of the nucleoplasmic localization of SR proteins. This is further supported by the observation of the cytoplasmic retention of both a SR34 mutant variant displaying R/S-to-G/T substitutions (SR34^GT^) (Stankovic *et al.,* 2016) and a SR45 variant displaying mutations within its RS2 (SR45mutRS2) (Fanara *et al.,* 2024). This suggests that RS-mutated variants are less effectively imported in the nucleus, which is in agreement with the absence of interaction of RS2ZmutRS variant with the plant transportin-SR (MOS14) thought to ensure the nuclear import of plant SR proteins (Xu *et al.,* 2011). Furthermore, the absence of interaction between the RS2ZmutRS variants and the SRPK kinases (SRPK3a, SRPK3b and SRPK4) also supports the observation of the cytoplasmic accumulation of these mutant variants, as the nuclear import of SR proteins by MOS14 requires phosphorylated residues (Lai *et al.,* 2000; Lai *et al.,* 2001). These SRPK kinases (SRPK3a, SRPK3b and SRPK4) might therefore contribute to the nuclear import of the RS2Z members conjointly with MOS14. This hypothesis is also supported by a recent study that linked an exclusive cytoplasmic localization of RS2Z32 to the down-regulation of phosphorylation levels of specific serine residues within the RS domain (but not within the SP domain) in an *srpk3 srpk4 srpk3b/srpk5* triple mutant (Wang *et al.,* 2023). All of these observations thus imply that nuclear import of RS2Z32 and RS2Z33 by the transportin-SR MOS14 requires prior phosphorylation of RS domain serine residues, likely by SRPK3a, SRPK3b and SRPK4. Interestingly, the RS2ZmutRRM+RS variant displays a fainter cytoplasmic fluorescent signal than the RS2ZmutRS, suggesting a diminution of the nuclear export of the RS2Z proteins, which is corroborated by the observed diminution of shuttling kinetics of this mutant variant compared to the native protein, mostly due to the alteration of the RNP2 motif.

### RS2Z proteins contact splicing partners through their RRM

dY2H results indicate that many residues of the RRM of RS2Z proteins (including the mutated residues Y14, Y46 and F48) are involved in the association to splicing factors, such as SR45, SCL30 and SR34. This corroborates a previous report stating that the RNA-binding domains of RS2Z33 (truncated protein harboring RRM+ZnKs only) were involved in the interactions with a second SCL subfamily member, namely SCL33 (Lopato *et al.,* 2002). Our dY2H and BiFC results additionally indicate the association of both RS2Z to SR45 involve both RS domains of SR45. Indeed, neither RS2Z32 nor RS2Z33 is able to interact with the SR45mutRS1+RS2 in yeast and plant cells **(Supplementary Figures S9 and S10)**. Reciprocally, the RS domain of RS2Z proteins also proved important to maintain this interaction in yeast cells **(Figure 4 and Supplementary Figure S7)**.

### RNA binding specificity of the RRM of RS2Z proteins

*Cis*-elements (or RNA motifs) bound by plant SR proteins have been studied through two complementary strategies: (i) *in silico* studies performed on RNA-Seq data of *sr* loss-of-function mutants searching for motifs common to differentially expressed genes, and (ii) SELEX experiments. Doing so, motifs associated to SCL30, SR45 and SR34 binding to RNAs were determined, all of which display purine-rich features (Yan *et al.,* 2017; Fanara *et al.,* 2024). In contrast, RS2Z proteins recognize a common *cis*-element displaying a pyrimidine-rich feature (U/C stretch) at the 3’ end of a central motif 5’-G[G/C]UA-3’. Interestingly, the sequences bound by RS2Z32 and RS2Z33 upstream and downstream of this central motif were more variable. These distinct specificities may stem from sequence divergence between the RRM of the two RS2Z proteins, e.g. the RS2Z32 Y62 residue is replaced by a histidine in RS2Z33, or the RS2Z32 A78 and N89 residues are replaced by phenylalanine residues capable of connecting RNA (Zeke *et al.,* 2022) in RS2Z33. However, these additional residues are not the main drivers of the RNA binding specificity of RS2Z proteins as mutations of three conserved residues in RNP2 and RNP1 motifs (Y14, Y46 and F48) were sufficient to disrupt the recovery of a significant motif with each RS2Z RRM. The enrichment in pyrimidine bases in an RNA motif is usually associated to the control of AS events (Ule *et al.,* 2006; Goers *et al.,* 2010; Uemura *et al.,* 2017). The binding of RS2Z33 to the *CALLOSE SYNTHASE5* (*CalS5*) pre-mRNA has been proposed to positively regulate the splicing removal of its intron 6 (Huang *et al.,* 2013). While the motif associated with both RS2Z32 and RS2Z33 (5’-CCGCUA-3’) is absent from the *CalS5* pre-mRNA sequence, the 5’-CUGCUA-3’ motif specifically bound by the RRM of RS2Z33 is present in both the exon 5 and the exon 27 of *CalS5*. The observation of a RS2Z33 binding motif in an exon upstream of the regulated intron 6 suggests that this SR protein might indeed regulate the splicing pattern of its putative target, *CalS5*, which supports the working hypothesis of Huang and colleagues (Huang *et al.,* 2013). The interactors of RS2Z33 (SR34, SR45 and SCL30) might not regulate this specific intron removal as their RNA binding sites (5’-AGGGAG-3’ for SR34 and 5’-AGAAGA-3’ for SR45 and SCL30) are present in relatively distant positions from the intron 6 (Yan *et al.,* 2017; Fanara *et al.,* 2024).

### Are RS2Z proteins playing redundant molecular functions?

While most of our results largely suggest a functional redundancy between RS2Z32 and RS2Z33 (e.g., identical expression profile and interactors, or similar dynamics and RNA motifs bound), our dY2H assays showed some noticeable differences between the RS2Z32 and RS2Z33 complexes. Specifically, the association of RS2Z32 to partners also contacted by RS2Z33 appeared to be weaker (e.g., Cyp59), which is demonstrated by a lighter blue staining and by the absence of yeast growth upon 100-times dilutions of the original culture. This was also visible in the domains involved in the interaction, as the RS2Z32 mutant variant displaying mutations to its RS (mutRS) was unable to associate to SR34, contrary to RS2Z33mutRS. Moreover, the nucleocytoplasmic shuttling activity of RS2Z32 was impaired upon RNP2 mutation, and the one of RS2Z33 was additionally affected upon ZnK2 alterations. All of these observations suggest diverse molecular mechanisms underlying the function of each of the RS2Z proteins.

The putative functional redundancy of the RS2Z proteins remains to be confirmed by complementation experiment of a double mutant. However, as a quintuple *rsz rs2z* mutant (Yan *et al.,* 2017) displays no phenotypic impairments, it is challenging to propose a specific biological function for the RS2Z members. One could speculate on the physiological role(s) of the RS2Z proteins by investigating the multiple stresses affecting the expression of their respective genes. Using available Arabidopsis RNA-seq resources (Zhang *et al.,* 2020; Yu *et al.,* 2022) (http://ipf.sustech.edu.cn/pub/athrna/) suggests that *RS2Z* genes are responsive to specific stresses for which they are distinctively transcriptionally regulated (e.g., dehydration, low red/far-red ratio or salt), sometimes in opposite directions (cold or methyl jasmonate) (Feng *et al.,* 2020), even though they can be co-regulated in some instances (e.g., dark treatments) (Pietzenuk *et al.,* 2016).

## Conclusion

In this study, we characterized the contribution of the RS, SP, RRM and two ZnKs domains, and their constitutive residues, to the function of the Arabidopsis RS2Z32 and RS2Z33 proteins investigating the many aspects defining a splicing factor. Serines of the RS domain control the nucleoplasmic diffusion of RS2Z32 and RS2Z33, as mutations applied to this sole domain led to a partial cytoplasmic retention combined to a nucleolar enrichment of these splicing factors. On the other hand, the RNA-binding domains of both RS2Z proteins (i.e. RNP2 and ZnK2) are involved in the shuttling activity of the RS2Z proteins, and the RRM itself is also involved in both protein-protein and protein-RNA interactions. The RRM specifically binds RNA molecules via the residues Y14, Y46 and F48, and the RNA motifs identified appear to be different from the generally observed purine-rich sequences recognized by SR45, SCL30 and SR34. Many charged residues in the RRM likely serve to contact splicing factors that also displays a shuttling dynamic (e.g., SR45 and SR34). A BiFC approach allowed the confirmation of several interactions observed in yeast cells. Among them, the connection between SR45, SCL30, SR34 or CypRS64 and the RS2Z proteins was visualized *in planta*. Collectively, our observations suggest that RS2Z proteins play highly redundant functions, which might explain the absence of phenotypes of a quintuple *rsz rs2z* mutant.

## Acknowledgments

We thank Dr Julien Spielmann, Dr Marine Joris, and Professor Moreno Galleni for helpful discussions. We thank Professor Frédéric Farnir for his advice on FLIP-shuttling statistical analyses.

## Author contributions

PM: conceptualization and directing the research; SF: conducting most experiments, with contributions of MS, SDF, and MV for SELEX experiments, and of MG for FLIP-shuttling assays; PM and SF: data analysis; PM and MH: supervision; SF: making the figures. SF, MH, and PM wrote the manuscript, with comments of all authors.

## Data availability

All data supporting the findings of the study are available within the paper and within its supplementary data published online.

## Supplementary data

**Supplementary Figure S1.** Scaled schemes depicting the native RS2Z32 and RS2Z33 proteins and their corresponding mutant variants. RA and AP variants correspond to the RS and SP domain, respectively, in which all the serines (17 in both genes for the RS domain, or 15/16 in RS2Z32/RS2Z33 for the SP domain) were substituted in alanines as shown in **Supplementary Table S5**. Critical aromatic amino acids (bold underlined) connecting the RNA in RNP2 and RNP1 motifs (Y14, Y46, F48) were substituted in alanines (Maris *et al.,* 2005; Califice *et al.,* 2012; Stankovic *et al.,* 2016). 1-284: RS2Z32 protein size in amino acids. 1-290: RS2Z33 protein size in amino acids.

**Supplementary Figure S2.** Toxicity and autoactivation assays of the native RS2Z32 and RS2Z33 and their corresponding mutant variants on a –Trp/X-a-gal medium. **(A)** Yeast transformants obtained using 100 ng of the following plasmids (clockwise): empty pGBTK7, pGBKT7:RS2Z, pGBKT7:RS2ZmutRRM, pGBKT7:RS2ZmutZnKs, pGBKT7:RS2ZmutRRM+ZnKs, pGBKT7:RS2ZmutRS, pGBKT7:RS2ZmutSP, and pGBKT7:RS2ZmutRS+SP. **(B)** Yeast transformants obtained using 100 ng of the following plasmids (clockwise): empty pGBTK7, pGBKT7:RS2ZmutRRM+RS, and pGBKT7:RS2ZmutRRM+SP. Similar results were obtained for pGADT7 constructs.

**Supplementary Figure S3.** Expression profiling of *RS2Z* genes in Arabidopsis organs. Quantitative RT-PCR analyses of *RS2Z* genes expression in Arabidopsis vegetative and reproductive organs. Values represent means ± SEM (from three biological replicates, each consisting of pools of organs from three plants) and are relative to *AT1G58050*. Data were analyzed by two-way analysis of variance (ANOVA) followed by Bonferroni multiple comparison post-test. Statistically significant differences between means of all genes within one tissue are indicated by asterisks (**P<0.01, ***P<0.001). Statistically significant differences between means for one gene between tissues are indicated by letters (P<0.05). n.s., not significant.

**Supplementary Figure S4.** Reporter lines expressing *pRS2Z32:EGFP* and *pRS2Z33:EGFP*. Both promoters are active in the seed coat and radicle **(A/a**), embryo **(B/b)**, hypocotyl **(C/c)** and cotyledons **(D/d),** two-week-old root epidermis **(E/e)** including root hairs **(F/f)**, leaf epidermis **(G/g-H/h)** including trichomes **(G/g)** and stomata [**(H/h)**, arrows], gynoecium (valves, stigma and style) **(I/i)**, androecium (anther and filament) **(J/j)**, pollen grains **(K/k)**, ovules **(L/l)** and funiculi [**(L/l)**, arrow)], and petals and sepals **(M/m)**. Red signals represent chlorophyll autofluorescence. At least three independent T3 homozygous lines were generated and analyzed, depicting similar fluorescence profiles.

**Supplementary Figure S5.** Neither RNP motifs nor ZnK domains control the nuclear distribution of RS2Z proteins. Subcellular fluorescence distribution in transient expression assays in tobacco leaf cells upon N-terminal EGFP-tagging of the mentioned mutant variants. Scale bars = 10 µm. Red signals represent chlorophyll autofluorescence. At least three independent transient events were generated and analyzed, depicting similar fluorescence profiles.

**Supplementary Figure S6.** Nucleocytoplasmic shuttling of mutant variants altered in the RRM, ZnKs or RRM+ZnKs.. FLIP-shuttling was assessed without (−LMB) or with LMB (+LMB) treatment in tobacco epidermal leaf cells. One hundred percent fluorescence indicates prebleach fluorescence intensity. Insets show the overlay of wild-type and mutant curves. Values are means ± SEM for at least 10 nuclei. A significant inhibitory effect of LMB is indicated by asterisks (*P < 0.05, **P < 0.01, ***P < 0.001, ****P < 0.0001). n.s., not significant. Three given time points [10s, 65s (50% of the time scale) or 129s] were statistically analyzed.

**Supplementary Figure S7.** Interactions of RS2Z32 and RS2Z33 with kinases, splicing-associated factors, and nuclear import factor upon serial dilutions. From the initial culture, dilutions to an OD600 of 0.25, 0.025 and 0.0025 were spotted on -Trp/-Leu/-His/X-α-Gal/AurA agar plates. Companion to **Figure 4**.

**Supplementary Figure S8.** Full-length RS2Z proteins strongly interact with only few Arabidopsis SR proteins. RS2Z32 **(A)** and RS2Z33 **(B)** interactions with RSZ, SC35 and SR subfamilies (mammalian orthologs), and RS2Z, SCL, RS and SR45 subfamilies (plant specific). **(C)** Interactions established between SR45 isoforms and truncated RS2Z32 or RS2Z33 variants lacking both RS and SP domains (ΔRS+SP), or the SP domain (ΔSP), or all domains but the RRM (ΔZnKs+RS+SP). From the initial culture, dilutions to an OD600 of 0.25 were spotted on - Trp/-Leu/-His/X-a-Gal/AurA agar plates. Positive interactions were confirmed by growth and blue staining. The absence of interaction is characterized at most by a shadow of dead cells.

**Supplementary Figure S9.** RS2Z proteins associate to SR45, partly through its RS1 domain. **(A)** Observed RS2Z interactions with SR45 mutant variants in yeast cells. From the mated culture, dilutions to an OD600 of 0.5 were spotted on -Trp/-Leu/-His/X-a-Gal/AurA agar plates. Positive interactions were confirmed by growth and blue staining. Yellow and purple squares are indicative of a weakened or abolished interaction, respectively, upon substitutions within the corresponding SR45 domain. A weakened interaction is characterized by the ability of several single colonies to grow, while an abolished interaction is characterized at most by a white halo of dead cells. **(B)** Interpretive summary of the interactions of RS2Z32 and RS2Z33 with SR45 mutant variants. Signs: +, positive interaction; -, absence of interaction; ±, weakened interaction.

**Supplementary Figure S10.** RS2Z proteins do not interact with the SR45mutRS1+RS2 mutant variant *in planta*. Bimolecular fluorescence complementation (BiFC) through co-expression of RS2Z32 or RS2Z33 fused to the C-terminal half of YFP (RS2Z32:^C^YFP or RS2Z33:^C^YFP) and SR45mutRS1+RS2 (SR45mutRS1+RS2:^N^YFP). Representative images of fluorescence reveal the absence of interaction of RS2Z32 or RS2Z33 with SR45mutRS1+RS2 in transient expression assay in tobacco leaf cells. At least three independent transient events were generated and analyzed, depicting similar fluorescence profiles. Scale bars = 10 µm. Red signals represent chlorophyll autofluorescence.

**Supplementary Figure S11.** The RRM and zink-knuckles of the RS2Z proteins do not conjointly provide a specific RNA motif binding. The non-significant RNA motifs were identified through 4 rounds of SELEX selection with the native RRM and the two ZnKs of RS2Z32 **(A)** and RS2Z33 **(B)**. The statistical significance (*E*-value) is indicated at the bottom of each consensus. Logos were redesigned using WebLogo (Crooks *et al*., 2004). All RNA motifs were discovered using the MEME tool (version 5.5.1) with 30 sequences.

**Supplementary Tables S1-S9.** List of primers used in this study, subdivided into experimental categories, and statistical analyses of FLIP-shuttling assays.

**Supplementary Tables S10-S15.** List of sequences submitted to MEME to identify motif consensus(es) for RNA-binding sites of the native RS2Z RRMs and the corresponding mutated RRMs or RRM+ZnKs recombinant proteins.

## References

Aubol, B. E., K. L. Hailey, L. Fattet, P. A. Jennings and J. A. Adams (2017). Redirecting SR Protein Nuclear Trafficking through an Allosteric Platform. Journal of Molecular Biology 429(14): 2178–2191.

Aubol, B. E., G. Wu, M. M. Keshwani, M. Movassat, L. Fattet, K. J. Hertel, X. D. Fu and J. A. Adams (2016). Release of SR Proteins from CLK1 by SRPK1: A Symbiotic Kinase System for Phosphorylation Control of Pre-mRNA Splicing. Molecular Cell 63(2): 218–228.

Bailey, T. L., M. Boden, F. A. Buske, M. Frith, C. E. Grant, L. Clementi, J. Ren, W. W. Li and W. S. Noble (2009). MEME SUITE: tools for motif discovery and searching. Nucleic Acids Research 37(Web Server issue): W202-208.

Barbosa Dos Santos, I. and S. W. Park (2019). Versatility of Cyclophilins in Plant Growth and Survival: A Case Study in *Arabidopsis*. Biomolecules 9(1).

Barta, A., M. Kalyna and A. S. Reddy (2010). Implementing a rational and consistent nomenclature for serine/arginine-rich protein splicing factors (SR proteins) in plants. The Plant Cell 22(9): 2926–2929.

Bender, J. and G. R. Fink (1994). AFC1, a LAMMER kinase from *Arabidopsis thaliana*, activates STE12-dependent processes in yeast. *Proceedings of the National Academy of Sciences*, USA 91(25): 12105–12109.

Bi, Y., Z. Deng, W. Ni, R. Shrestha, D. Savage, T. Hartwig, S. Patil, S. H. Hong, Z. Zhang, J. A. Oses-Prieto, K. H. Li, P. H. Quail, A. L. Burlingame, S. L. Xu and Z. Y. Wang (2021). Arabidopsis ACINUS is O-glycosylated and regulates transcription and alternative splicing of regulators of reproductive transitions. Nature Communications 12(1): 945.

Califice, S., D. Baurain, M. Hanikenne and P. Motte (2012). A single ancient origin for prototypical serine/arginine-rich splicing factors. Plant Physiology 158(2): 546–560.

Cavaloc, Y., C. F. Bourgeois, L. Kister and J. Stévenin (1999). The splicing factors 9G8 and SRp20 transactivate splicing through different and specific enhancers. RNA 5(3): 468–483.

Chen, M. and J. L. Manley (2009). Mechanisms of alternative splicing regulation: insights from molecular and genomics approaches. Nature Reviews Molecular Cell Biology 10(11): 741–754.

Cléry, A., M. Blatter and F. H. Allain (2008). RNA recognition motifs: boring? Not quite. Current Opinion in Structural Biology 18(3): 290–298.

Crooks, G. E., G. Hon, J. M. Chandonia and S. E. Brenner (2004). WebLogo: a sequence logo generator. Genome Research 14(6): 1188–1190.

Cruz, T. M., R. F. Carvalho, D. N. Richardson and P. Duque (2014). Abscisic acid (ABA) regulation of *Arabidopsis* SR protein gene expression. International Journal of Molecular Sciences 15(10): 17541–17564.

Curtis, M. D. and U. Grossniklaus (2003). A gateway cloning vector set for high-throughput functional analysis of genes in planta. Plant Physiology 133(2): 462–469.

Czechowski, T., M. Stitt, T. Altmann, M. K. Udvardi and W. R. Scheible (2005). Genome-wide identification and testing of superior reference genes for transcript normalization in Arabidopsis. Plant Physiology 139(1): 5–17.

Daubner, G. M., A. Cléry and F. H. Allain (2013). RRM-RNA recognition: NMR or crystallography…and new findings. Current Opinion in Structural Biology 23(1): 100–108.

De Franco, S., J. Vandenameele, A. Brans, O. Verlaine, K. Bendak, C. Damblon, A. Matagne, D. J. Segal, M. Galleni, J. P. Mackay and M. Vandevenne (2019). Exploring the suitability of RanBP2-type Zinc Fingers for RNA-binding protein design. Scientific Reports 9(1): 2484.

Drechsel, G., A. Kahles, A. K. Kesarwani, E. Stauffer, J. Behr, P. Drewe, G. Rätsch and A. Wachter (2013). Nonsense-mediated decay of alternative precursor mRNA splicing variants is a major determinant of the *Arabidopsis* steady state transcriptome. The Plant Cell 25(10): 3726–3742.

English, A. C., K. S. Patel and A. E. Loraine (2010). Prevalence of alternative splicing choices in *Arabidopsis thaliana*. BMC Plant Biology 10: 102.

Fanara, S., M. Schloesser, M. Joris, S. De Franco, M. Vandevenne, F. Kerff, M. Hanikenne and P. Motte (2024). The Arabidopsis SR45 splicing factor bridges the splicing mc fachinery and the exon-exon junction complex. J Exp Bot 75(8): 2280–2298.

Feng, G., M. J. Yoo, R. Davenport, J. L. Boatwright, J. Koh, S. Chen and W. B. Barbazuk (2020). Jasmonate induced alternative splicing responses in *Arabidopsis*. Plant Direct 4(8): e00245.

Goers, E. S., J. Purcell, R. B. Voelker, D. P. Gates and J. A. Berglund (2010). MBNL1 binds GC motifs embedded in pyrimidines to regulate alternative splicing. Nucleic Acids Research 38(7): 2467–2484.

Gullerova, M., A. Barta and Z. J. Lorković (2006). AtCyp59 is a multidomain cyclophilin from *Arabidopsis thaliana* that interacts with SR proteins and the C-terminal domain of the RNA polymerase II. RNA 12(4): 631–643.

Hanikenne, M., I. N. Talke, M. J. Haydon, C. Lanz, A. Nolte, P. Motte, J. Kroymann, D. Weigel and U. Krämer (2008). Evolution of metal hyperaccumulation required cis-regulatory changes and triplication of *HMA4*. Nature 453(7193): 391–395.

Haynes, C. and L. M. Iakoucheva (2006). Serine/arginine-rich splicing factors belong to a class of intrinsically disordered proteins. Nucleic Acids Research 34(1): 305–312

Huang, X. Y., J. Niu, M. X. Sun, J. Zhu, J. F. Gao, J. Yang, Q. Zhou and Z. N. Yang (2013). CYCLIN-DEPENDENT KINASE G1 is associated with the spliceosome to regulate *CALLOSE SYNTHASE5* splicing and pollen wall formation in *Arabidopsis*. The Plant Cell 25(2): 637–648.

Huang, Y. and J. A. Steitz (2001). Splicing factors SRp20 and 9G8 promote the nucleocytoplasmic export of mRNA. Molecular Cell 7(4): 899–905.

Jo, S. H., H. J. Park, A. Lee, H. Jung, J. M. Park, S. Y. Kwon, H. S. Kim, H. J. Lee, Y. S. Kim, C. Jung and H. S. Cho (2022). The Arabidopsis cyclophilin CYP18-1 facilitates PRP18 dephosphorylation and the splicing of introns retained under heat stress. The Plant Cell 34(6): 2383–2403.

Kalyna, M., S. Lopato and A. Barta (2003). Ectopic expression of atRSZ33 reveals its function in splicing and causes pleiotropic changes in development. Molecular Biology of the Cell 14(9): 3565–3577.

Kalyna, M., C. G. Simpson, N. H. Syed, D. Lewandowska, Y. Marquez, B. Kusenda, J. Marshall, J. Fuller, L. Cardle, J. McNicol, H. Q. Dinh, A. Barta and J. W. Brown (2012). Alternative splicing and nonsense-mediated decay modulate expression of important regulatory genes in *Arabidopsis*. Nucleic Acids Research 40(6): 2454–2469.

Kashkan, I., K. Timofeyenko and K. Růžička (2022). How alternative splicing changes the properties of plant proteins. *Quantitative Plant Biology* 3(e14): 1–11.

Kataoka, N., J. L. Bachorik and G. Dreyfuss (1999). Transportin-SR, a nuclear import receptor for SR proteins. Journal of Cell Biology 145(6): 1145–1152.

Kim, D. Y., M. Scalf, L. M. Smith and R. D. Vierstra (2013). Advanced proteomic analyses yield a deep catalog of ubiquitylation targets in *Arabidopsis*. The Plant Cell 25(5): 1523–1540.

Königs, V., C. de Oliveira Freitas Machado, B. Arnold, N. Blümel, A. Solovyeva, S. Löbbert, M. Schafranek, I. Ruiz De Los Mozos, I. Wittig, F. McNicoll, M. H. Schulz and M. Müller-McNicoll (2020). SRSF7 maintains its homeostasis through the expression of Split-ORFs and nuclear body assembly. Nature Structural & Molecular Biology 27(3): 260–273.

Lai, M. C., R. I. Lin, S. Y. Huang, C. W. Tsai and W. Y. Tarn (2000). A human importin-beta family protein, transportin-SR2, interacts with the phosphorylated RS domain of SR proteins. Journal of Biological Chemistry 275(11): 7950–7957.

Lai, M. C., R. I. Lin and W. Y. Tarn (2001). Transportin-SR2 mediates nuclear import of phosphorylated SR proteins. *Proceedings of the National Academy of Sciences*, USA 98(18): 10154–10159.

Liang, Q., Q. Geng, L. Jiang, M. Liang, L. Li, C. Zhang and W. Wang (2019). Protein methylome analysis in *Arabidopsis* reveals regulation in RNA-related processes. Journal of Proteomics 213: 103601.

Long, J. C. and J. F. Cáceres (2009). The SR protein family of splicing factors: master regulators of gene expression. Biochemical Journal 417(1): 15–27.

Lopato, S., C. Forstner, M. Kalyna, J. Hilscher, U. Langhammer, K. Indrapichate, Z. J. Lorković and A. Barta (2002). Network of interactions of a novel plant-specific Arg/Ser-rich protein, atRSZ33, with atSC35-like splicing factors. Journal of Biological Chemistry **277**(42): 39989-39998.

Lorković, Z. J. and A. Barta (2004). Compartmentalization of the splicing machinery in plant cell nuclei. Trends in Plant Science 9(12): 565–568.

Lorković, Z.J., Lopato, S., Pexa, M., Lehner, R., and Barta, A. (2004). Interactions of *Arabidopsis* RS domain containing cyclophilins with SR proteins and U1 and U11 small nuclear ribonucleoprotein-specific proteins suggest their involvement in pre-mRNA Splicing. Journal of Biological Chemistry 279: 33890–33898.

Lorković, Z.J., Hilscher, J., and Barta, A. (2008). Co-localisation studies of *Arabidopsis* SR splicing factors reveal different types of speckles in plant cell nuclei. Experimental Cell Research 314: 3175–3186.

Manley, J. L. and A. R. Krainer (2010). A rational nomenclature for serine/arginine-rich protein splicing factors (SR proteins). Genes & Development 24(11): 1073–1074.

Manuel, J. M., N. Guilloy, I. Khatir, X. Roucou and B. Laurent (2023). Re-evaluating the impact of alternative RNA splicing on proteomic diversity. Frontiers in Genetics 14: 1089053.

Maris, C., C. Dominguez and F. H. Allain (2005). The RNA recognition motif, a plastic RNA- binding platform to regulate post-transcriptional gene expression. The FEBS Journal 272(9): 2118–2131.

Marquez, Y., M. Höpfler, Z. Ayatollahi, A. Barta and M. Kalyna (2015). Unmasking alternative splicing inside protein-coding exons defines exitrons and their role in proteome plasticity. Genome Research 25(7): 995–1007.

Meyer, K., Koester, T., and Staiger, D. (2015). Pre-mRNA Splicing in Plants: *In Vivo* Functions of RNA-Binding Proteins Implicated in the Splicing Process. Biomolecules 5: 1717–1740.

Ni, X., A. C. Joerger, A. Chaikuad and S. Knapp (2023). Structural insights into a regulatory mechanism of FIR RRM1-FUSE interaction. Open Biology 13(5): 230031.

Pabis, M., G. M. Popowicz, R. Stehle, D. Fernández-Ramos, S. Asami, L. Warner, S. M. García-Mauriño, A. Schlundt, M. L. Martínez-Chantar, I. Díaz-Moreno and M. Sattler (2019). HuR biological function involves RRM3-mediated dimerization and RNA binding by all three RRMs. Nucleic Acids Research 47(2): 1011–1029.

Palusa, S. G., G. S. Ali and A. S. Reddy (2007). Alternative splicing of pre-mRNAs of Arabidopsis serine/arginine-rich proteins: regulation by hormones and stresses. The Plant Journal 49(6): 1091–1107.

Palusa, S. G. and A. S. Reddy (2010). Extensive coupling of alternative splicing of pre-mRNAs of serine/arginine (SR) genes with nonsense-mediated decay. New Phytologist 185(1): 83–89.

Pfaffl, M. W. (2001). A new mathematical model for relative quantification in real-time RT-PCR. Nucleic Acids Research 29(9): e45.

Pietzenuk, B., C. Markus, H. Gaubert, N. Bagwan, A. Merotto, E. Bucher and A. Pecinka (2016). Recurrent evolution of heat-responsiveness in *Brassicaceae COPIA* elements. Genome Biology 17(1): 209.

Qüesta, J. I., J. Song, N. Geraldo, H. An and C. Dean (2016). *Arabidopsis* transcriptional repressor VAL1 triggers Polycomb silencing at *FLC* during vernalization. Science 353(6298): 485–488.

Rausin, G., V. Tillemans, N. Stankovic, M. Hanikenne and P. Motte (2010). Dynamic nucleocytoplasmic shuttling of an Arabidopsis SR splicing factor: role of the RNA-binding domains. Plant Physiology 153(1): 273–284.

Remy, E., T. R. Cabrito, P. Baster, R. A. Batista, M. C. Teixeira, J. Friml, I. Sá-Correia and P. Duque (2013). A major facilitator superfamily transporter plays a dual role in polar auxin transport and drought stress tolerance in Arabidopsis. The Plant Cell 25(3): 901–926.

Spielmann, J., H. Ahmadi, M. Scheepers, M. Weber, S. Nitsche, M. Carnol, B. Bosman, J. Kroymann, P. Motte, S. Clemens and M. Hanikenne (2020). The two copies of the zinc and cadmium ZIP6 transporter of *Arabidopsis halleri* have distinct effects on cadmium tolerance. Plant, Cell & Environment 43(9): 2143–2157.

Stankovic, N., M. Schloesser, M. Joris, E. Sauvage, M. Hanikenne and P. Motte (2016). Dynamic Distribution and Interaction of the Arabidopsis SRSF1 Subfamily Splicing Factors. Plant Physiology 170(2): 1000–1013.

Talke, I. N., M. Hanikenne and U. Krämer (2006). Zinc-dependent global transcriptional control, transcriptional deregulation, and higher gene copy number for genes in metal homeostasis of the hyperaccumulator *Arabidopsis halleri*. Plant Physiology 142(1): 148–167.

Thompson, H. L., W. Shen, R. Matus, M. Kakkar, C. Jones, D. Dolan, S. Grellscheid, X. Yang, N. Zhang, S. Mozaffari-Jovin, C. Chen, X. Zhang, J. F. Topping and K. Lindsey (2023). MERISTEM-DEFECTIVE regulates the balance between stemness and differentiation in the root meristem through RNA splicing control. Development 150(7).

Tillemans, V., L. Dispa, C. Remacle, M. Collinge and P. Motte (2005). Functional distribution and dynamics of Arabidopsis SR splicing factors in living plant cells. The Plant Journal 41(4): 567–582.

Tillemans, V., I. Leponce, G. Rausin, L. Dispa and P. Motte (2006). Insights into nuclear organization in plants as revealed by the dynamic distribution of Arabidopsis SR splicing factors. The Plant Cell 18(11): 3218–3234.

Uemura, Y., T. Oshima, M. Yamamoto, C. J. Reyes, P. H. Costa Cruz, T. Shibuya and Y. Kawahara (2017). Matrin3 binds directly to intronic pyrimidine-rich sequences and controls alternative splicing. Genes to Cells 22(9): 785–798.

Ule, J., G. Stefani, A. Mele, M. Ruggiu, X. Wang, B. Taneri, T. Gaasterland, B. J. Blencowe and R. B. Darnell (2006). An RNA map predicting Nova-dependent splicing regulation. Nature 444(7119): 580–586.

Wang, T., X. Wang, H. Wang, C. Yu, C. Xiao, Y. Zhao, H. Han, S. Zhao, Q. Shao, J. Zhu, Y. Zhao, P. Wang and C. Ma (2023). *Arabidopsis* SRPKII family proteins regulate flowering via phosphorylation of SR proteins and effects on gene expression and alternative splicing. New Phytologist 238(5): 1889–1907.

Wu, Z., D. Zhu, X. Lin, J. Miao, L. Gu, X. Deng, Q. Yang, K. Sun, D. Zhu, X. Cao, T. Tsuge, C. Dean, T. Aoyama, H. Gu and L. J. Qu (2016). RNA Binding Proteins RZ-1B and RZ-1C Play Critical Roles in Regulating Pre-mRNA Splicing and Gene Expression during Development in Arabidopsis. The Plant Cell 28(1): 55–73.

Xu, S., Z. Zhang, B. Jing, P. Gannon, J. Ding, F. Xu, X. Li and Y. Zhang (2011). Transportin-SR is required for proper splicing of *Resistance* genes and plant immunity. PLoS Genetics 7(6): e1002159.

Yan, Q., X. Xia, Z. Sun and Y. Fang (2017). Depletion of *Arabidopsis* SC35 and SC35-like serine/arginine-rich proteins affects the transcription and splicing of a subset of genes. PLoS Genetics 13(3): e1006663.

Yang, Q., M. Coseno, G. M. Gilmartin and S. Doublié (2011). Crystal structure of a human cleavage factor CFI(m)25/CFI(m)68/RNA complex provides an insight into poly(A) site recognition and RNA looping. Structure 19(3): 368–377.

Yu, Y., H. Zhang, Y. Long, Y. Shu and J. Zhai (2022). Plant Public RNA-seq Database: a comprehensive online database for expression analysis of ∼45 000 plant public RNA-Seq libraries. Plant Biotechnology Journal 20(5): 806–808.

Zeke, A., E. Schád, T. Horváth, R. Abukhairan, B. Szabó and A. Tantos (2022). Deep structural insights into RNA-binding disordered protein regions. Wiley Interdisciplinary Reviews: RNA 13(5): e1714.

Zhang, H., F. Zhang, Y. Yu, L. Feng, J. Jia, B. Liu, B. Li, H. Guo and J. Zhai (2020). A Comprehensive Online Database for Exploring approximately 20,000 Public *Arabidopsis* RNA-Seq Libraries. Molecular Plant 13(9): 1231–1233.

Zhang, X. N. and S. M. Mount (2009). Two alternatively spliced isoforms of the Arabidopsis SR45 protein have distinct roles during normal plant development. Plant Physiology 150(3): 1450–1458.

