## Supplementary Figures for "The Arabidopsis RS2Z32 and RS2Z33 proteins are dynamic splicing factors whose RNA recognition motif (RRM) domain contributes to protein-protein and protein-RNA interactions"

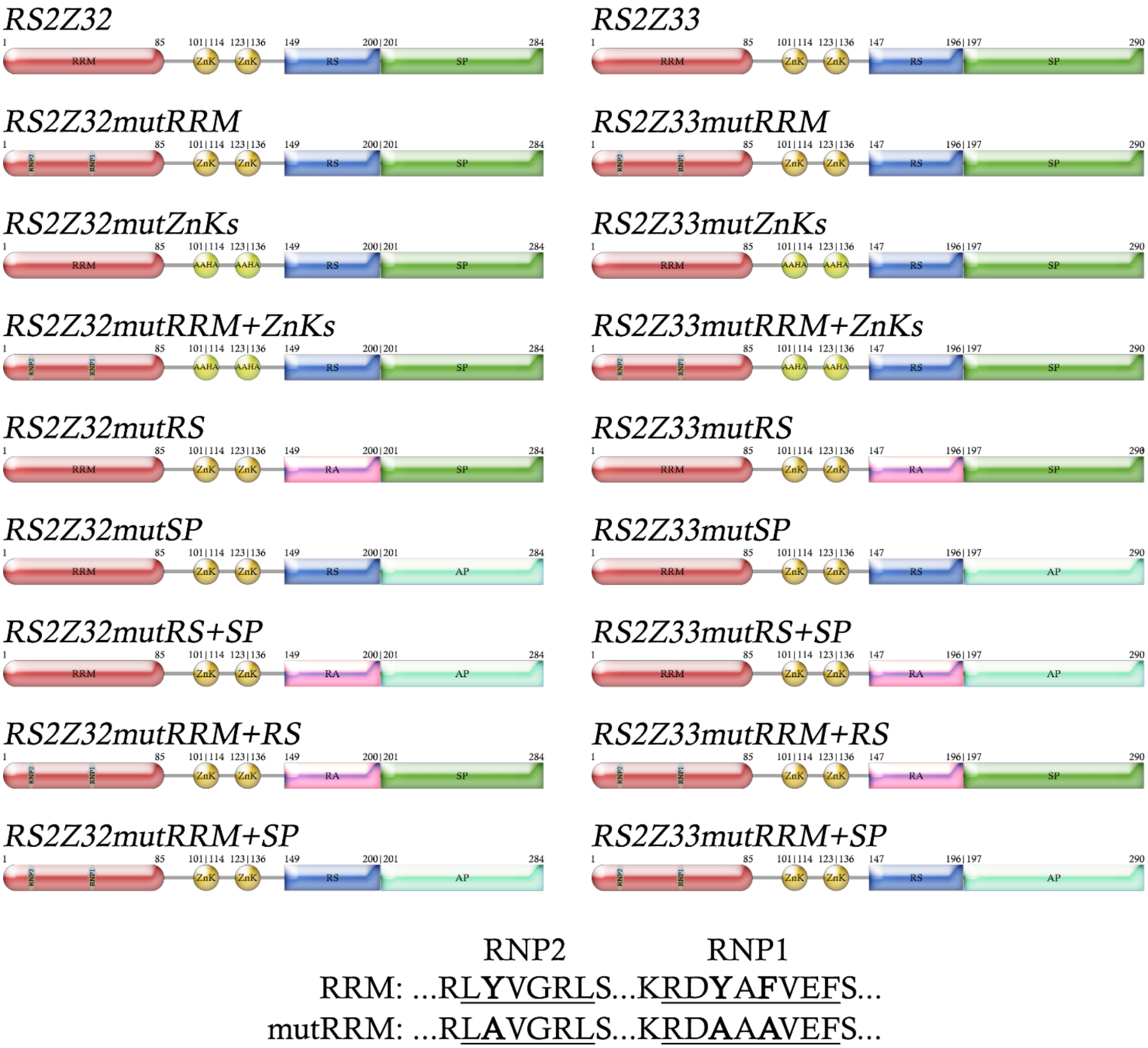


Supplementary Figure S1

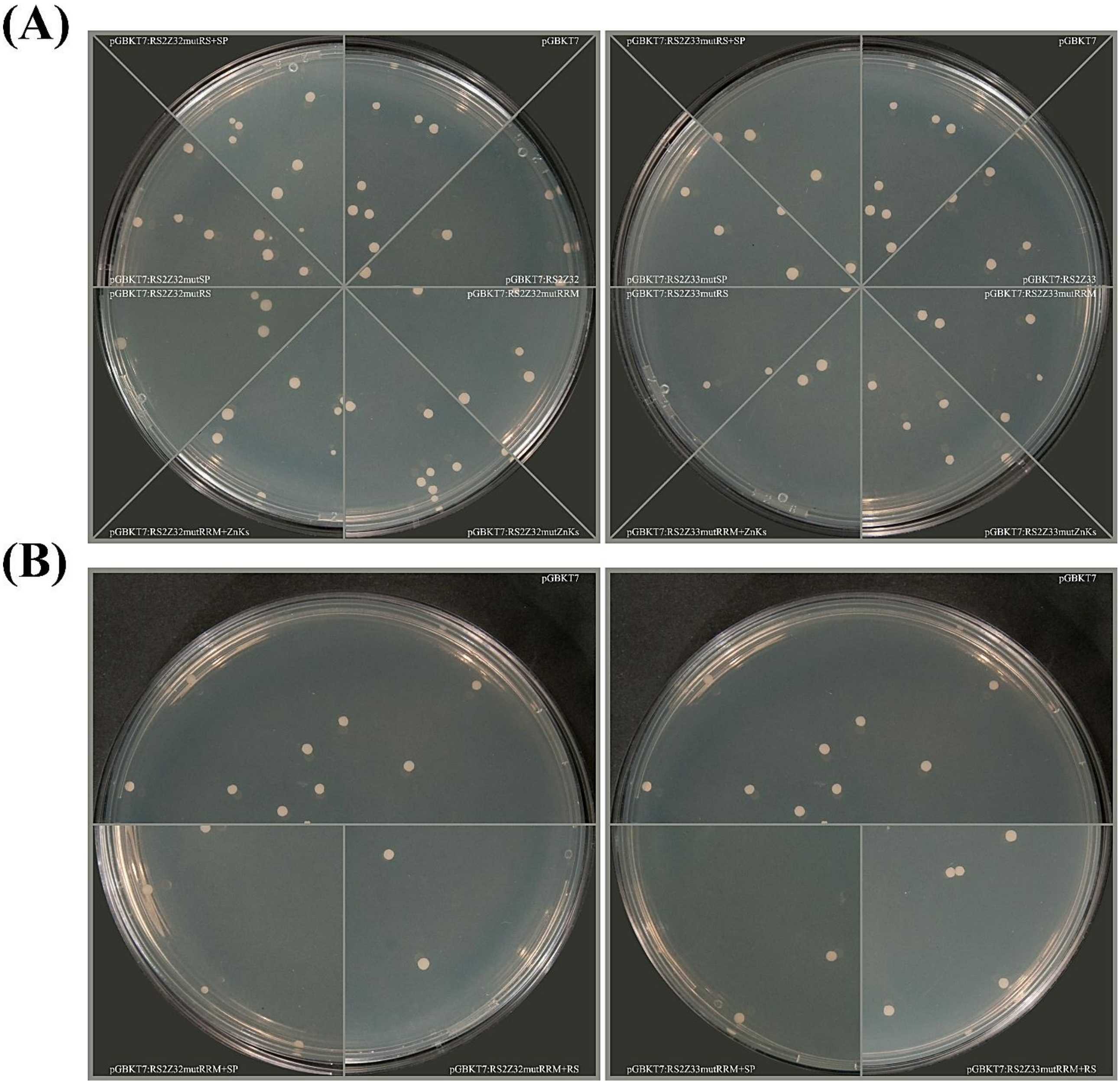


Supplementary Figure S2

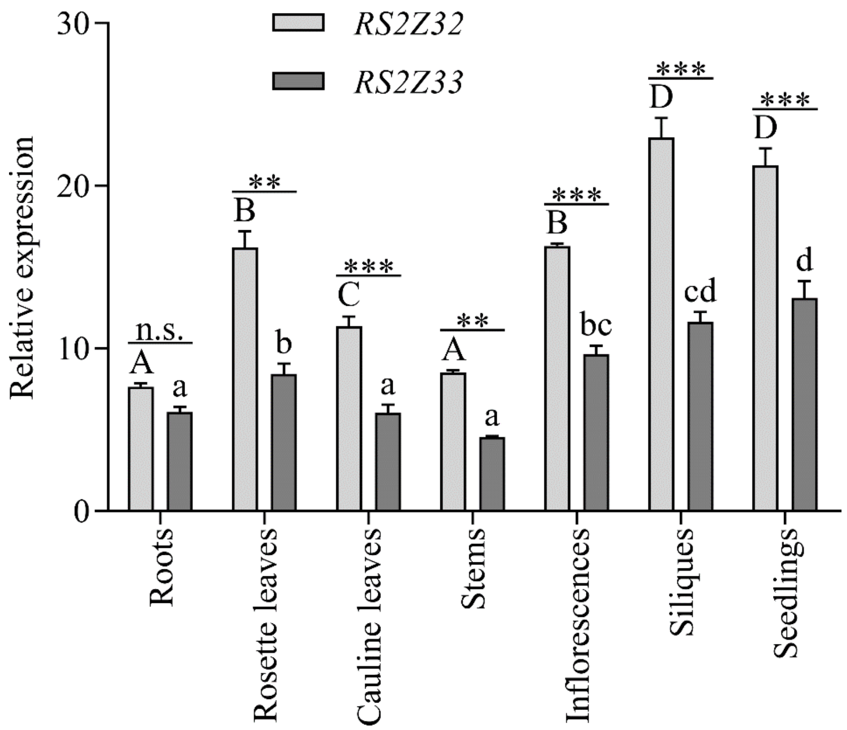


Supplementary Figure S3

*pRS2Z32:EGFP*

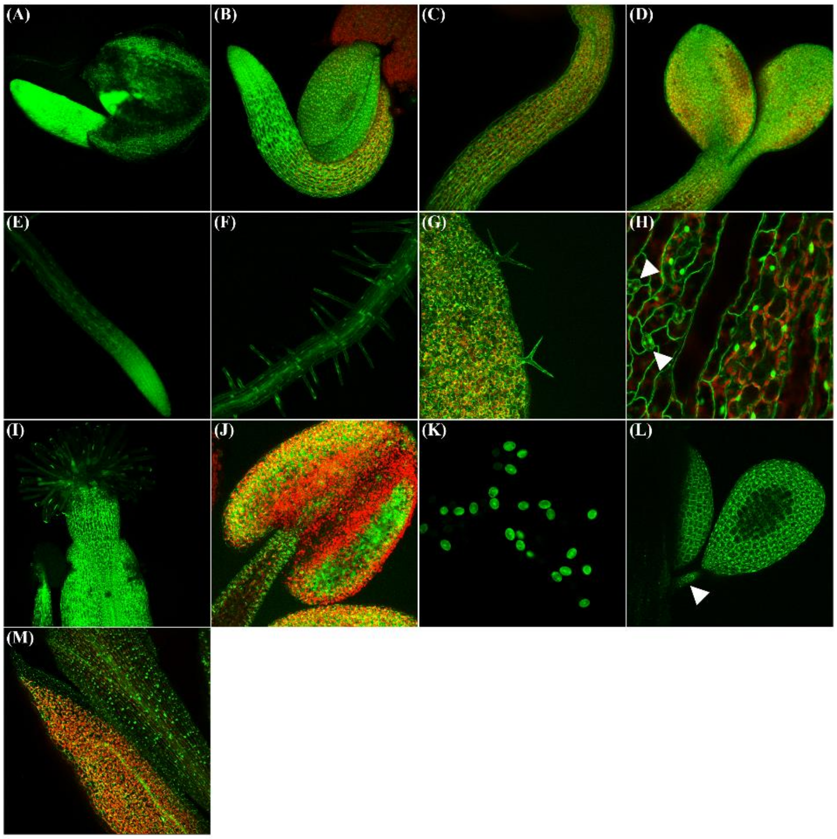


*pRS2Z33:EGFP*

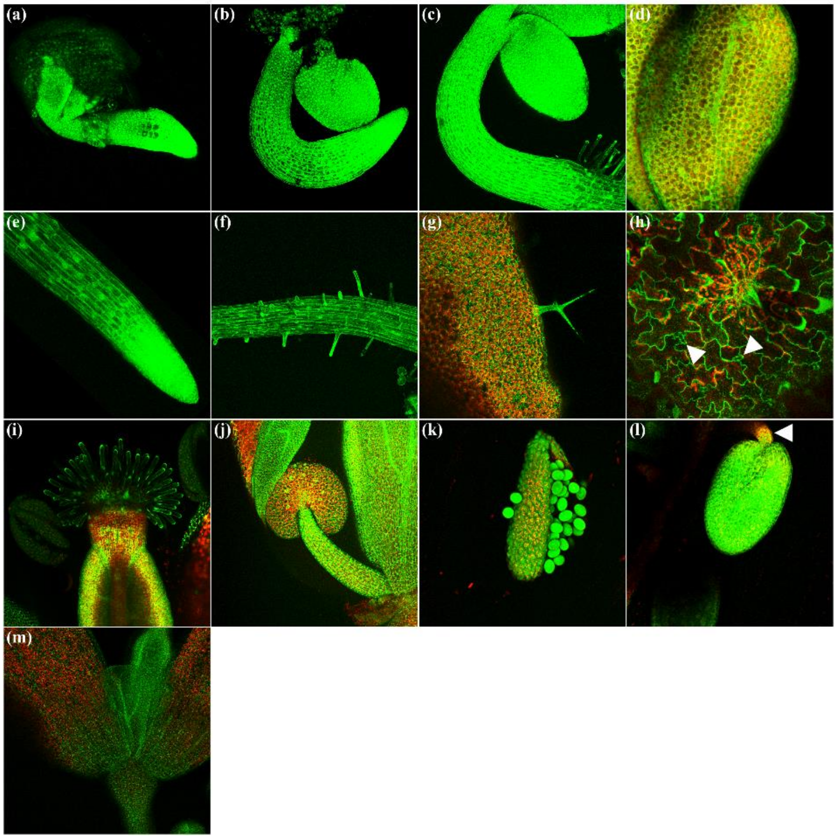


Supplementary Figure S4

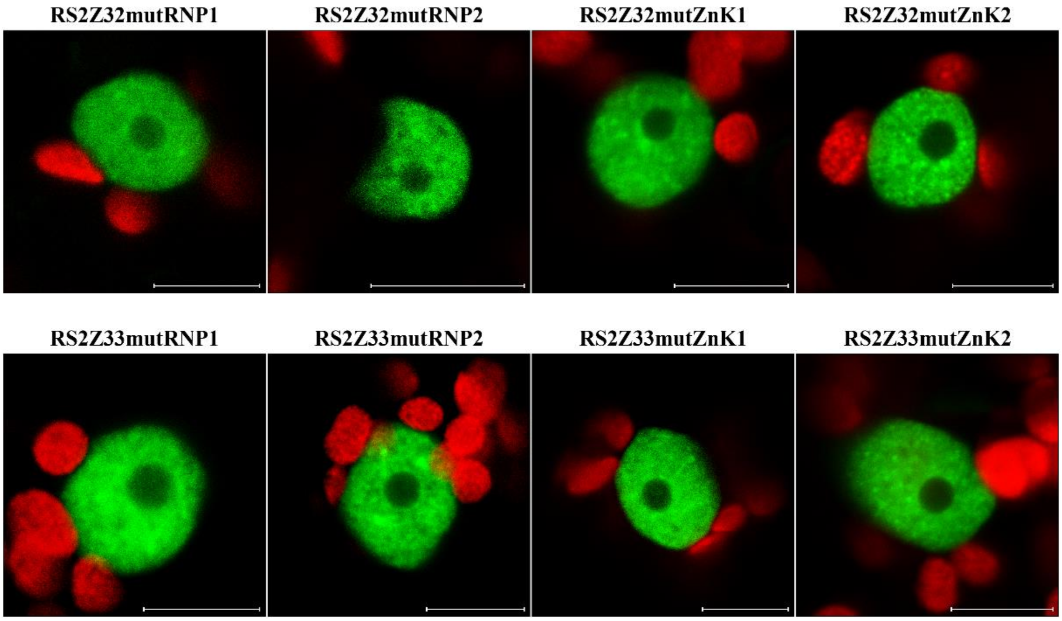


Supplementary Figure S5

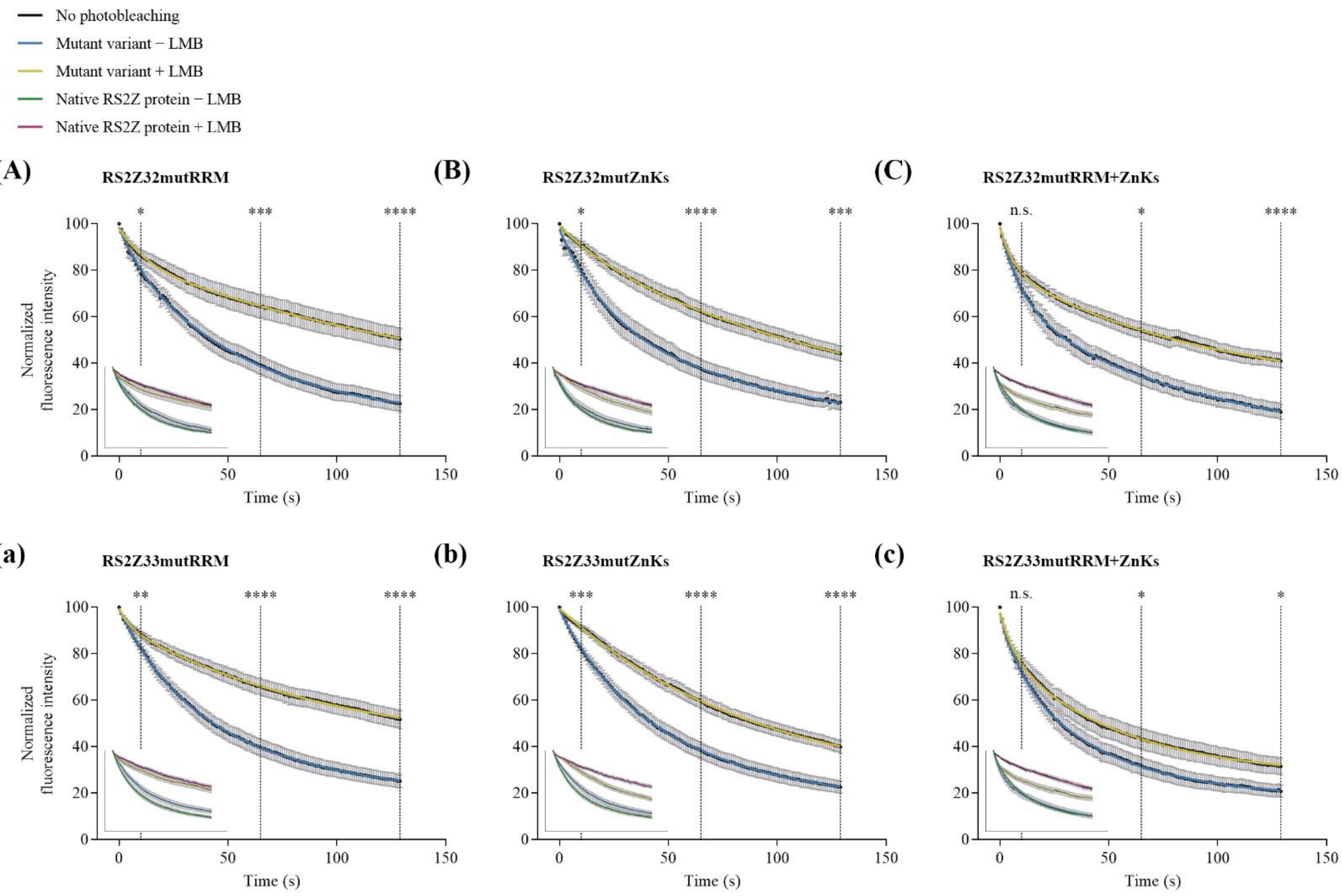


Supplementary Figure S6

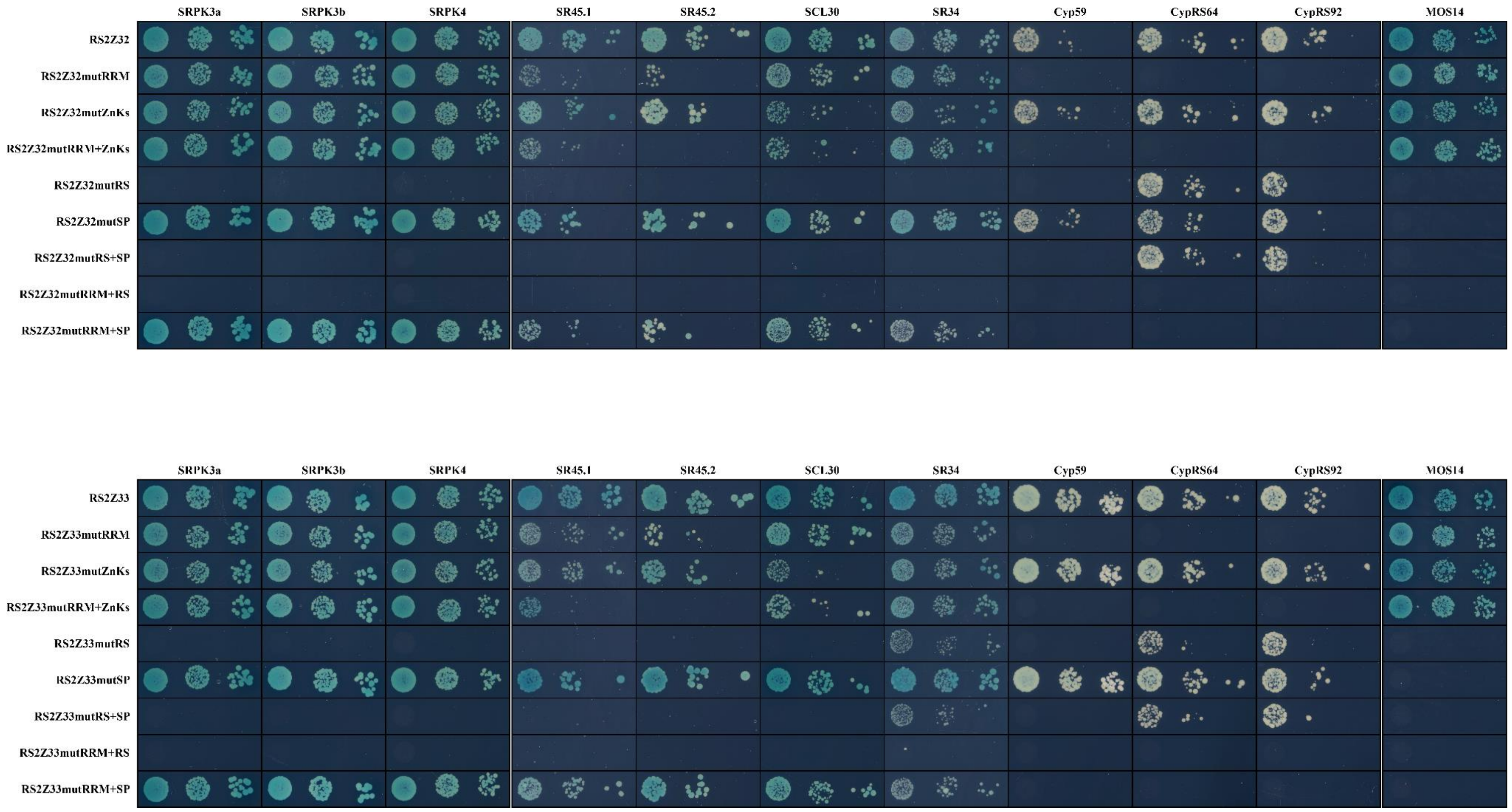


Supplementary Figure S7

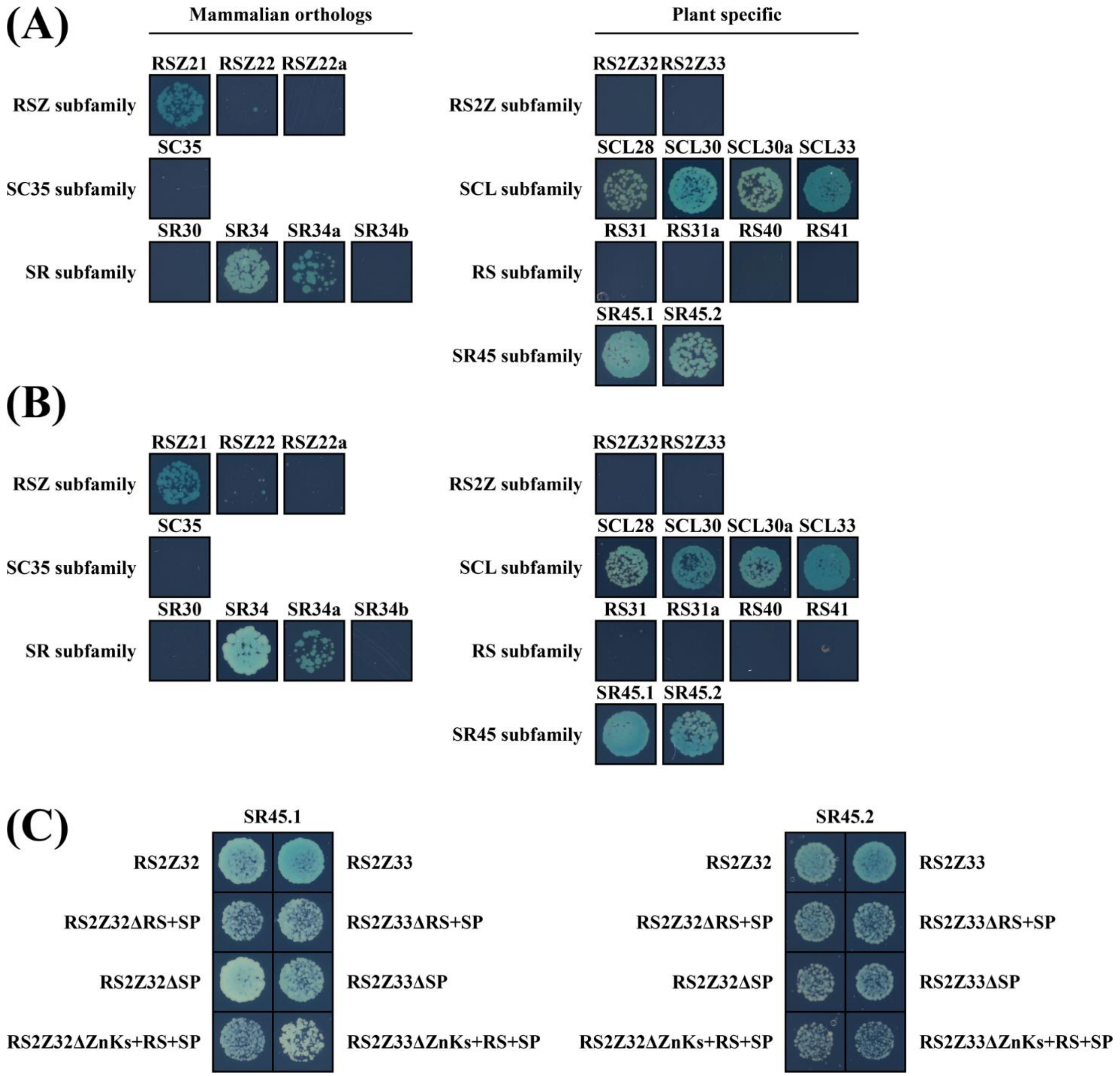


Supplementary Figure S8

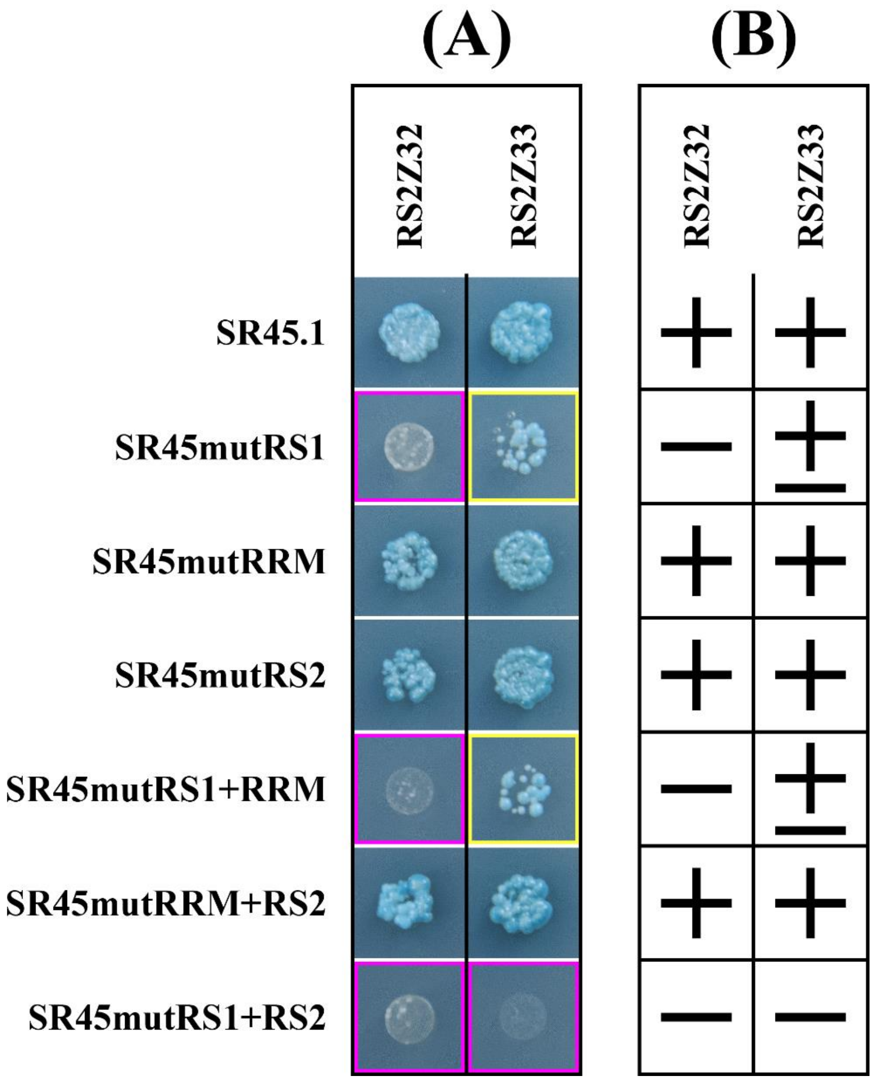


Supplementary Figure S9

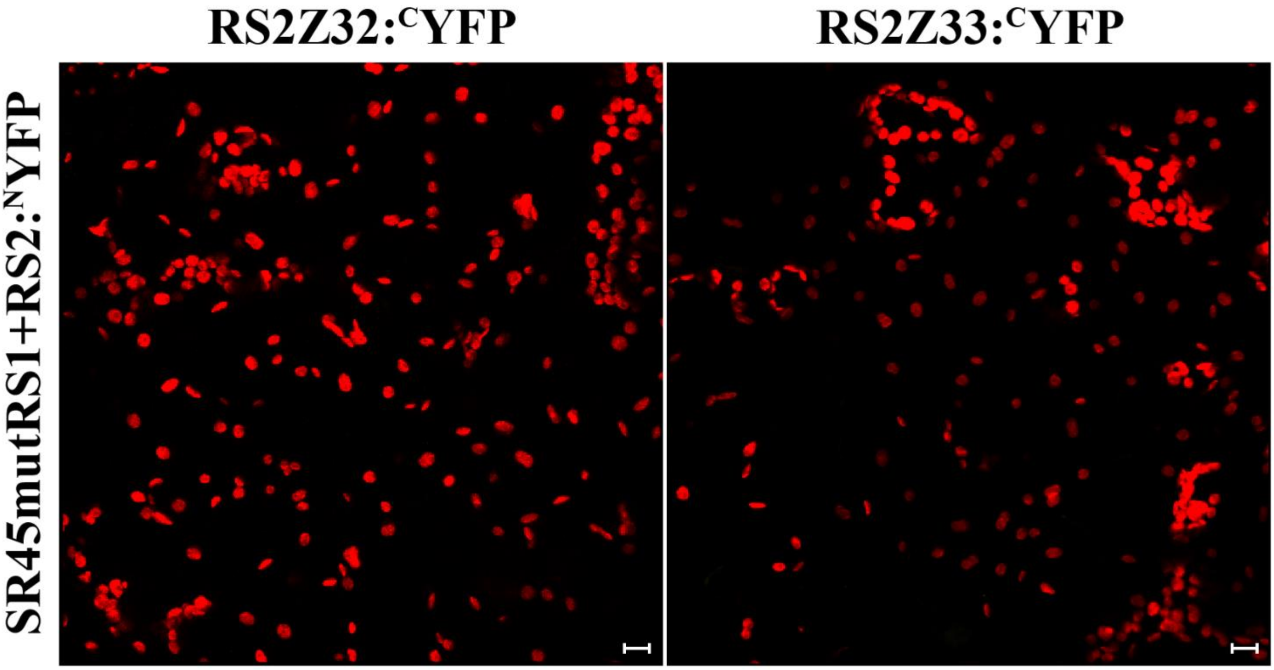


Supplementary Figure S10

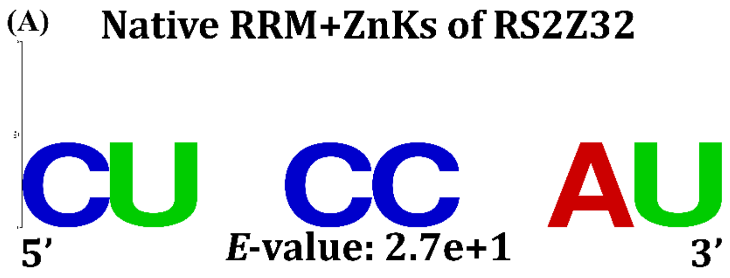


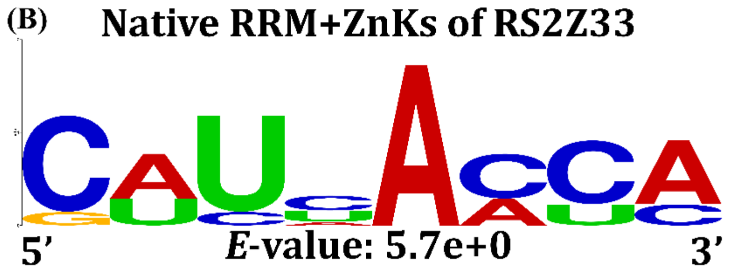


Supplementary Figure S11
