## Supplementary Tables for "The Arabidopsis RS2Z32 and RS2Z33 proteins are dynamic splicing factors whose RNA recognition motif (RRM) domain contributes to protein-protein and protein-RNA interactions"

| Target | Forward (5' – 3') | Reverse (5' – 3') | Construct | Literature |
|---|---|---|---|---|
| **SR proteins** | | | | |
| RS2Z32 | GGGAATTCATGCCTCGCTATGATGATCGC | GGGGATCCTCAAGGTGACTCACTGCCTTTAGG | pGADT7 / pGBKT7 | (Fanara *et al.*, 2024) |
| RS2Z32ΔSP | GGGAATTCATGCCTCGCTATGATGATCGC | GGGGATCCTCATGGACTGCGTGATCT | pGADT7 / pGBKT7 | This study |
| RS2Z32ΔRS+SP | GGGAATTCATGCCTCGCTATGATGATCGC | GGGGATCCTCATCCACCCTGCCTGGC | pGADT7 / pGBKT7 | This study |
| RS2Z32ΔZnKs+RS+SP | GGGAATTCATGCCTCGCTATGATGATCGC | GGGGATCCTCAGCGACCAGAACCAGGAG | pGADT7 / pGBKT7 | This study |
| RS2Z33 | GGGAATTCATGCCTCGCTATGATGATCGC | GGATCCAGGAGACTCACTTCCTCTAGG | pGADT7 / pGBKT7 | (Fanara *et al.*, 2024) |
| RS2Z33ΔSP | GGGAATTCATGCCTCGCTATGATGATCGC | GGGGATCCTCATGGACTGCGTGATCT | pGADT7 / pGBKT7 | This study |
| RS2Z33ΔRS+SP | GGGAATTCATGCCTCGCTATGATGATCGC | GGGGATCCTCATCCACTGCGCCTAAGCT | pGADT7 / pGBKT7 | This study |
| RS2Z33ΔZnKs+RS+SP | GGGAATTCATGCCTCGCTATGATGATCGC | GGGGATCCTCAGCGACCAGCTCCAGG | pGADT7 / pGBKT7 | This study |
| RS31 | GGGAATTCATGAGGCCAGTGTTCGTCG | GGGGATCCTCAAGGTCTTCCTCTTGGGACT | pGADT7 | (Fanara *et al.*, 2024) |
| RS31a | GGGAATTCATGAGACATGTGTACGTTGGGAATT | GGGGATCCTCAACCTCTTGCTCTTTGAATCG | pGADT7 / pGBKT7 | (Fanara *et al.*, 2024) |
| RS40 | GGGAATTCATGAAGCCAGTCTTCTGTGGG | GGGGATCCTCACTCGTCAGCTGGTGGC | pGBKT7 | (Fanara *et al.*, 2024) |
| RS41 | GGGAATTCATGAAGCCTGTCTTTTGCGG | GGGGATCCTCATTCCTCTGCTGGCGG | pGADT7 / pGBKT7 | (Fanara *et al.*, 2024) |
| RSZ21 | GGCCCGGG**T**ATGACGAGGGTTTATGTCGGG | GGGGATCCTCACACCCCATTGGCATATG | pGADT7 | (Fanara *et al.*, 2024) |
| RSZ22 | GAATTCATGTCACGTGTGTACGTCG | GGATCCTCAGCTCCTGCTTCTGC | pGADT7 / pGBKT7 | (Fanara *et al.*, 2024) |
| RSZ22a | GGGAATTCATGTCGCGTGTGTATGTTGGTAAT | GGCCCGGGTCAGCTCCGGCTTCTGC | pGADT7 / pGBKT7 | (Fanara *et al.*, 2024) |
| SC35 | GGGAATTCATGTCGCACTTCGGAAGGTC | GGGGATCCTCATTCCGCAGCATAAGGAGA | pGADT7 / pGBKT7 | (Fanara *et al.*, 2024) |
| SCL28 | GGGAATTCATGGCTAGAGCGAGAAGCCG | GGGGATCCTCAACGACTTAAGGATCGAGAACG | pGBKT7 | (Fanara *et al.*, 2024) |
| SCL30 | GGGAATTCATGAGGAGATACAGTCCGCCTTATTA | GGCTGCAGTCATCTTGGAGATACCTCCACAGAC | pGBKT7 | (Fanara *et al.*, 2024) |
| SCL30 | GGGAATTCATGAGGAGATACAGTCCGCCTTATTA | GGGAGCTCTCATCTTGGAGATACCTCCACAGAC | pGADT7 | (Fanara *et al.*, 2024) |
| SCL30a | GGGAATTCATGAGAGGAAGGAGCTACACGC | GGCCCGGGTCACTGGCTTGGAGAACGG | pGADT7 / pGBKT7 | (Fanara *et al.*, 2024) |
| SCL33 | GGGAATTCATGAGGGGAAGGAGCTACACTCC | GGGGATCCTCACTGGCTTGGTGAACGG | pGADT7 / pGBKT7 | (Fanara *et al.*, 2024) |
| SR30 | GGGGCCATGG**GG**ATGAGTGGGCGATTTTCTCGGTC | GGATCCACCAGATATCACAGGTGAAAC | pGADT7 / pGBKT7 | (Stankovic et al., 2016) |
| SR34 | GAATTCATGAGCAGTCGTTCGA | AGGGATCCCCTCGATGGACTC | pGADT7 / pGBKT7 | (Stankovic et al., 2016) |
| SR34a | GGGCCATGG**GG**ATGAGTGGGCGATTTTCTCGGTC | AGGGATCCCACACTGCCTTCGC | pGADT7 / pGBKT7 | (Stankovic et al., 2016) |
| SR34b | GAATTCATGAGCAGCCGTTCG | GGATCCTATCAATGCGATCCAATG | pGBKT7 | (Stankovic et al., 2016) |
| SR45.1 / SR45.2 | CGGGATCC**GG**ATGGCGAAACCAAGTCGTGGC | CCGAGCTCTTAAGTTTTACGAGGTGGAGGTGGTGG | pGADT7 | (Stankovic et al., 2016) |
| SR45.1 / SR45.2 | CGGGATCC**GG**ATGGCGAAACCAAGTCGTGGC | GGCTGCAGAGTTTTACGAGGTGGAGG | pGBKT7 | (Stankovic et al., 2016) |

| | | | | |
|---|---|---|---|---|
| SR45mut(RRM+)RS2 | GGGGCGCGCCATGGCGAAACCAAGTCGTG | GGGGATCCAGTTTTACGAGGTGGAGGTGGTGGTGG | pGADT7(+) / pGBKT7(+) | (Fanara *et al.*, 2024) |
| SR45mutRS1(+RRM+RS2) | GGGGCGCGCCATGGCGAAACCAGCTCGT | GGGGATCCAGTTTTACGAGGTGGAGGTGGTGGTGG | pGADT7(+) / pGBKT7(+) | (Fanara *et al.*, 2024) |
| **hnRNP-like** | | | | |
| GRP7 | GGGAATTCATGGCGTCCGGTGATGTTG | GGGGATCCTTACCATCCTCCACCACCACC | pGADT7 | (Fanara *et al.*, 2024) |
| GRP8 | GGGAATTCATGTCTGAAGTTGAGTACCGGTGC | GGGGATCCTTACCAGCCGCCACCAC | pGADT7 / pGBKT7 | (Fanara *et al.*, 2024) |
| RZ-1B | CCGAATTCATGAAAGATAGAGAAAACGATGGAAATC | GGGGATCCCTACCAACGTTCATATGATGAAGGTC | pGBKT7 | (Fanara *et al.*, 2024) |
| RZ-1C | CCGAATTCATGGCTGCAAAAGAAGGTAGTAGG | GGGGATCCTTAATAACGGTCAAAAGTGGACG | pGBKT7 | (Fanara *et al.*, 2024) |
| **PRPs** | | | | |
| PRP38 | GGGGATCCTTATGGCAAACAGAACAGATCCG | GGCTCGAGGTCCCTGAGGGGTTTCATTCC | pGADT7 | (Fanara *et al.*, 2024) |
| **Kinases** | | | | |
| AFC1 | GGCCCGGGTATGCAAAGCAGTGTGTATCGTGATAA | GGGGATCCTTAGTTCTTTTGGTTGTATAAAAATGGATGAG | pGADT7 | (Fanara *et al.*, 2024) |
| AFC2 | GGCATATGATGGAGATGGAGCGTGTGC | GGGGATCCCTATCTTCTCCTTGCGAAAAACG | pGADT7 / pGBKT7 | (Fanara *et al.*, 2024) |
| AFC3 | GGGAATTCATGATAGCTAACGGATTCGAGAGTATG | GGGGATCCTCAACTTGAGCTCTTAAAGAAAGGATG | pGADT7 | (Fanara *et al.*, 2024) |
| SRPK1 | GGGAATTCATGTCTTGTTCATCTTCTTCCGGAT | GGGGATCCTCACCCTTTGATATGCAAGTTGTTC | pGADT7 | (Fanara *et al.*, 2024) |
| SRPK2 | GGGAATTCATGTCGTGTTCATCCTCATCTGG | GGCCCGGGTCAAGAACATGAACCTTTGATCTGC | pGADT7 | (Fanara *et al.*, 2024) |
| SRPK3a | GGGAATTCATGGCGGATGAGAAGAACGG | GGGGATCCTTAAGTACGAAGATCACGAGCTGATTG | pGADT7 / pGBKT7 | (Fanara *et al.*, 2024) |
| SRPK3b/SRPK5 | GGGAATTCATGGCGGAGGACAAAAACAAC | GGGGATCCTCACTTCGGTTCAGAGACATCAATAG | pGADT7 | (Fanara *et al.*, 2024) |
| SRPK4 | GGGAATTCATGGAGGCGGAGAAGTGGAACAG | CGGGATCCAATTGCTAGCTTAAGAGTGGAGGAGCTT | pGADT7 / pGBKT7 | (Stankovic et al., 2016) |
| **Cyclophilines** | | | | |
| Cyp59 | GGGAATTCATGTCAGTTCTTATTGTGACGAGCC | GGGGATCCTCATCTATCCCTTCTCTCATGTCTAGCT | pGADT7 | (Fanara *et al.*, 2024) |
| CypRS64 | CGGAATTCATGACTAAAAAGAAGAATCCTAATGTTTT | TATAGAGCTCTCAATCCGCATAGCTAACCAG | pGADT7 | (Stankovic et al., 2016) |
| Cyp65 | GGGAATTCATGGGGAAGAAACAACACAGCA | GGGGATCCTTACCAGCTAGAGAAATCTTTAAACCCTG | pGADT7 | (Fanara *et al.*, 2024) |
| Cyp71 | GGGAATTCATGGAGGAAGAATCTAAGAATGGCG | GGCCCGGGCTAAGATTTCGGAACGGTGACATTAA | pGADT7 | (Fanara *et al.*, 2024) |
| CypRS92 | GGGAATTCATGGCAAAAAAGAAGAATCCACAGG | GGCCCGGGTTAATCATAGGCTACTAATCCCTTTTTCCC | pGADT7 | (Fanara *et al.*, 2024) |
| **EJC(-related)** | | | | |
| ACINUS(RRM) | CCGAATTCATGCTTCCTGCTAATGATCAAGAAGC | CCGGATCCTCACTTGTTATTATTCGCTGCAAGTTTAG | pGADT7 / pGBKT7 | (Fanara *et al.*, 2024) |
| ALY4 | CCGAATTCATGTCTGGAGCATTGAATATGACTCTTG | CCGGATCCTCAAGAGGTGTTCATGGCATCAGC | pGADT7 / pGBKT7 | (Fanara *et al.*, 2024) |
| eIF4A3 | CCGAATTCATGGCGACAGCGAATCCTGG | CCGGATCCTCAGATAAGATCAGCTACATTCATTGGC | pGADT7 / pGBKT7 | (Fanara *et al.*, 2024) |
| MAGO | GGGAATTCATGGCCGCGGAAGAAGC | GGGGATCCCTAGATAGGCTTGATTTTGAAGTGCAG | pGADT7 / pGBKT7 | (Fanara *et al.*, 2024) |

| | | | | |
|---|---|---|---|---|
| PININ | GGCATATGATGGGAGACACCGCCTTG | CCGAATTCTTAGAGAACCTCATGTTTAATATCTTCC | pGADT7 / pGBKT7 | (Fanara *et al.*, 2024) |
| SAP18 | CCGAATTCATGGCTGAAGCAGCGAGAAGAC | CCGGATCCTCAGTAAATTGCCACATCCAGATAATC | pGBKT7 | (Fanara *et al.*, 2024) |
| Y14 | CCGAATTCATGGCGAACATAGAATCAGAAGCAGTC | CCGGATCCTCAGTAACGTCTTCTCGGACTTCTTG | pGADT7 / pGBKT7 | (Fanara *et al.*, 2024) |
| **MOSes** | | | | |
| MOS12 | GGCATATGATGATTTACACTGCTATCGACAATTTTTAC | GGGGATCCTTAATGGTGCCTACGACGGTCT | pGBKT7 | (Fanara *et al.*, 2024) |
| MOS14 | GGCCCGGG**T**ATGGAGCATCAGAACGCGG | GGGGATCCTGATACAGGAGCAGTAACCAGATTCA | pGADT7 | (Fanara *et al.*, 2024) |

**Supplementary Table S1:** Primers used to amplify coding sequences of potential interactors used in directed yeast two-hybrid assays. Restriction sites used are underlined. Bold blue bases are used to preserve the open reading frame in final construction. Cells colored in light green or light orange represent, respectively, interactors identify through yeast two-hybrid screens using either a commercially available Arabidopsis cDNA library (Mate and Plate Library-Universal Arabidopsis, Clontech) or a custom cDNA library (Make Your Own "Mate & Plate" Library System, Clontech).

| Expression profiling and protein localization | | | |
|---|---|---|---|
| **Target** | **Forward (5' – 3')** | **Reverse (5' – 3')** | **Construct** ***Remark*** |
| pRS2Z32 | GGCCTGCAGGTCTTTAATGGGCTTTAGAGTTGATATAAGG | CCGGTACCTGTCAAGCTGCAAAATATTTT | pMDC32:pRS2Z32:RS2Z32:EGFP<br>pMDC32:pRS2Z32:EGFP |
| RS2Z32 | CGGGCGCGCCATGCCTCGCTATGAT | CCTTAATTAAAGGTGACTCACTGCCTTTAGG | pMDC32:pRS2Z32:RS2Z32:EGFP |
| pRS2Z33 | GGCCTGCAGGAATAATAAAGTCATAATTTATGAA | CCGGTACCTGTCAAGCTACAAAATCAAAAA | pMDC32:pRS2Z33:RS2Z33:EGFP<br>pMDC32:pRS2Z33:EGFP |
| RS2Z33 | TTGGCGCGCCATGCCTCGCTATGATGATCGCTATG | CCTTAATTAAAGGAGACTCACTTCCTCTAGGGGAAGTG | pMDC32:pRS2Z33:RS2Z33:EGFP |
| EGFP | CCTTAATTAA**C**ATGGTGAGCAAGGGCGAGGAG | CCGAGCTCTTACTTGTACAGCTCGTCCATGC | pMDC32:pRS2Z32:RS2Z32:EGFP<br>pMDC32:pRS2Z33:RS2Z33:EGFP |
| EGFP | CCGGCGCGCC**C**ATGGTGAGCAAGGGCGAGGAG | CCGAGCTCTTACTTGTACAGCTCGTCCATGC | pMDC32:pRS2Z32:EGFP<br>pMDC32:pRS2Z33:EGFP |

**Supplementary Table S2:** Primers used for constructs for expression profiling and protein localization *in planta*. Restriction sites used are underlined. Bold blue bases are used to ensure the open reading frame in final construction.

| PCR-based point mutagenesis | | | |
|---|---|---|---|
| **Gene** | **Forward (5’ – 3’)** | **Reverse (5’ – 3’)** | **Mutant** |
| **RS2Z32** | GATGTGGATATGAAGCGTGATGCTGCCGCTGTTGAATTTAGTGATCCTCG | CGAGGATCACTAAATTCAACAGCGGCAGCATCACGCTTCATATCCACATC | RS2Z32mutRNP1 |
| **RS2Z32** | TGGAAACACTCGCCTCGCTGTTGGTCGCTTATCA | TGATAAGCGACCAACAGCGAGGCGAGTGTTTCCA | RS2Z32mutRNP2 |
| **RS2Z32** | GGTTCTGGTCGCGCTTTTAATGCTGGTGTCGATGGC | GCCATCGACACCAGCATTAAAAGCGCGACCAGAACC | RS2Z32mutZnK1.1 |
| **RS2Z32** | ACTGGGCCCGAGACGCCACAGCAGGAGACT | AGTCTCCTGCTGTGGCGTCTCGGGCCCAGT | RS2Z32mutZnK1.2 |
| **RS2Z32** | ACTGGAAGAATAAAGCTTACCGCGCTGGTGAAAGAGGACA | TGTCCTCTTTCACCAGCGCGGTAAGCTTTATTCTTCCAGT | RS2Z32mutZnK2.1 |
| **RS2Z32** | AGAGGACACATTGAGAGAAACGCCAAAAACAGTCCTAGTCCAAA | TTTGGACTAGGACTGTTTTTGGCGTTTCTCTCAATGTGTCCTCT | RS2Z32mutZnK2.2 |
| **RS2Z33** | TGGATATGAAGCGAGATGCTGCTGCCGTTGAATTTGGTGATCCCC | GGGGATCACCAAATTCAACGGCAGCAGCATCTCGCTTCATATCCA | RS2Z33mutRNP1 |
| **RS2Z33** | GCTATGGGAACACTCGTCTTGCCGTTGGCCGATTATCATCGA | TCGATGATAATCGGCCAACGGCAAGACGAGTGTTCCCATAGC | RS2Z33mutRNP2 |
| **RS2Z33** | GAGCTGGTCGCGCTTTTAACGCTGGTGTAGATGG | CCATCTACACCAGCGTTAAAAGCGCGACCAGCTC | RS2Z33mutZnK1.1 |
| **RS2Z33** | ATTGGGCTCGTGACGCCACAGCAGGGGACT | AGTCCCCTGCTGTGGCGTCACGAGCCCAAT | RS2Z33mutZnK1.2 |
| **RS2Z33** | TGGAAGAACAAGGCTTACCGTGCTGGAGAGAGAGGACACA | TGTGTCCTCTCTCTCCAGCACGGTAAGCCTTGTTCTTCCA | RS2Z33mutZnK2.1 |
| **RS2Z33** | AGGACACATTGAGAGAAACGCCAAAAACCAGCCCAAGAAG | CTTGGGCTGGTTTTTGGCGTTTCTCTCAATGTGTCCT | RS2Z33mutZnK2.2 |

**Supplementary Table S3:** Primers used for PCR-based point mutagenesis in the RRM and in ZnK1/ZnK2 domains of *RS2Z32* and *RS2Z33*. Critical aromatic amino acids connecting the RNA in RNP2 and RNP1 motifs were substituted in alanine (Maris *et al.,* 2005; Califice *et al.,* 2012; Stankovic *et al.,* 2016). The substituting codons are underlined in the sequence.

| Primers | Sequence (5' – 3') | |
|---|---|---|
| | RS2Z32 | RS2Z33 |
| 1 | GGGAATTCATGCCTCGCTATGATGATCGC | GGGAATTCATGCCTCGCTATGATGATCGC |
| 2 | GGGGATCCTCAAGGTGACTCACTGCCTTTAGG | GGATCCAGGAGACTCACTTCCTCTAGG |
| 3 | GGGGATCCTCAAGGTGCCTCAGCGCC | GGGGATCCTTAAGGAGCCTCAGCTCCTCTAGG |
| 4 | GGCGCTGAGGCACCTTGAGGATCC | CCTAGAGGAGCTGAGGCTCCTTAAGGATCC |
| 5 | CGGGCCCAGTGGCCATCGACACCACAATTAAAACAGCG | ATCCCGAGAACCACGAGGTGCCCCTCGTGAAAACTCCACAGT |
| 6 | TGGCCACTGGGCCCG | CACCTCGTGGTTCTCGGGAT |
| 7 | TCAAGGTGCCTCAGCGCC | TTAAGGAGCCTCAGCTCCTCTAGG |
| 8 | GGACAGAGCACGCGCTCCTAAGGCAATGGAGCGATCTGTATC | GGAGGAGAGATCACGCAGTCCAAAGCGGATGGATGACGCTCTA |
| 9 | GGAGGACAGATCACGCAGTCCTAAGGCAATGGAGCGAGCTGTA | GGAGGACAGATCACGCAGTCCTAAGGCAATGGAGCGAGCTGTA |

**Supplementary Table S4:** Primers used to construct *RS2Z32* and *RS2Z33* mutant variants. Restriction sites used (*Eco*RI et *Bam*HI) are underlined.

| **Synthetic genes** | |
|---|---|
| **Mutant variant** | **Coding sequence** |
| RS+SP domains to RA+AP domains of *RS2Z32*<br><br>(411 bp, 32 S-to-A substitutions) | GGA**GCA**TAT**GCC**AGG**GCA**CCAGTCAAA**GCC**CGC**GCC**CCTCGTCGCCGAAGG**GCA**CCA**GCA**CGT**GCA**CGT**GCT**TAC**GCT**CGAGGT<br>CGC**GCA**TAC**GCT**CGA**GCC**CGA**GCC**CCAGTGAGAAGAGAGAAA**GCA**GTGGAGGACAGA**GCA**CGC**GCT**CCTAAGGCAATGGAGCGA<br>**GCT**GTA**GCT**CCCAAAGGTAGGGACCAA**GCA**CTG**GCT**CCAGACCGAAAAGTGATAGATGCA**GCA**CCAAAGCGTGGA**GCA**GACTAT<br>GATGGT**GCA**CCAAAAGAGAATGGTAATGGCAGGAAC**GCT**GCG**GCT**CCCATTGTTGGAGGTGGTGAA**GCT**CCTGTTGGACTTAAT<br>GGTCAAGACAGG**GCA**CCGATTGATGATGAGGCTGAGCTT**GCA**CGTCCT**GCC**CCTAAAGGC**GCT**GAG**GCA**CCTTGA |
| RS+SP domains to RA+AP domains of *RS2Z33*<br><br>(435 bp, 33 S-to-A substitutions) | GGA**GCA**TAC**GCC**AGG**GCA**CCTGTAAGA**GCC**CGT**GCT**CCTCGTCGTAGAAGA**GCA**CCA**GCA**CGG**GCT**CTT**GCA**CGT**GCA**CGA**GCA**<br>TAC**GCA**CGA**GCA**CGA**GCC**CCGGTGAGAAGAAGAGAGAGG**GCT**GTGGAGGAGAGA**GCA**CGC**GCT**CCAAAGCGGATGGATGAC**GCT**<br>CTA**GCC**CCAAGAGCCAGAGATCGT**GCT**CCGGTTCTTGATGATGAAGGC**GCC**CCAAAGATCATAGACGGG**GCA**CCACCACCA**GCA**<br>CCAAAGCTTCAAAAGGAAGTCGGA**GCT**GACCGTGACGGTGGT**GCC**CCCCAAGACAATGGCAGAAAC**GCT**GTTGTC**GCT**CCTGTT<br>GTAGGAGCCGGTGGTGAC**GCTGCC**AAAGAGGACCGG**GCA**CCTGTTGATGATGATTACGAGCCAAACCGCACT**GCC**CCTAGAGGA<br>**GCT**GAG**GCT**CCTTAA |

**<u>Supplementary Table S5:</u>** Synthetic genes provided by GenScript where serine codons within RS and SP domains of *RS2Z32* and *RS2Z33* genes were substituted with alanine codons (red and bold).

| BiFC | | | |
|---|---|---|---|
| **Target** | **Forward (5' – 3')** | **Reverse (5' – 3')** | **Construct** |
| RS2Z32 | CGGGATCCATGCCTCGCTATGAT | GTGGTACCAGGTGACTCACTGCCTT | pBI121:35S:RS2Z32:$^{N}$YFP<br>pBI121:35S:RS2Z32:$^{C}$YFP |
| RS2Z33 | CGGGATCCATGCCTCGCTATGAT | GTGGTACCAGGAGACTCACTTCCTC | pBI121:35S:RS2Z33:$^{N}$YFP<br>pBI121:35S:RS2Z33:$^{C}$YFP |
| CypRS64 | GGGGCGCGCCATGACTAAAAAGAAGAATCCTAATG | GGTTAATTAAATCCGCATAGCTAACCAG | pBI121:35S:CypRS64:$^{N}$YFP |

**Supplementary Table S6:** Primers used to confirm protein-protein interactions *in planta* (BiFC experiments). Restriction sites used are underlined. Bold blue bases are used to ensure the open reading frame in final construction.

| SELEX | |
|---|---|
| **Synthesis of the DNA initial library** | |
| **Randomized initial library** | ***T7* promoter** |
| TCCCGCTCGTCGTCT*NNNNNNNNNNNNNNNNNNNNNNNNN*CCGCATCGTCCTCCCT | GAAATTAATACGACTCACTATAGGGAGGACGATGCGG |

| Recombinant gene construction | | | |
|---|---|---|---|
| **Target** | **Forward (5' – 3')** | **Reverse (5' – 3')** | **Construct** |
| RS2Z32 (native & mutated RRM) | GGGGATCCATGCCTCGCTATGATGATCGC | GGGAATTCTCAGCGACCAGAACCAGGAG | pGEX6P1 |
| RS2Z32 (native RRM+ZnKs) | GGGGATCCATGCCTCGCTATGATGATCGC | GGGAATTCTCATCCACCCTGCCTGGC | pGEX6P1 |
| RS2Z33 (native & mutated RRM) | GGGGATCCATGCCTCGCTATGATGATCGC | GGGAATTCTCAGCGACCAGCTCCAGG | pGEX6P1 |
| RS2Z33 (native RRM+ZnKs) | GGGGATCCATGCCTCGCTATGATGATCGC | GGGAATTCTCATCCACTGCGCCTAAGCT | pGEX6P1 |

**Supplementary Table S7:** Primers used for SELEX experiments (De Franco *et al.*, 2019). Red bases correspond to the *T7* promoter. Restriction sites used are underlined.

| Statistical analyses of FLIP-shuttling assays | | | | | | | | | |
|---|---|---|---|---|---|---|---|---|---|
| **Comparisons** | | **Time** | | | | | | **Fluorescence intensities at 50% of the time scale (65s)** | |
| Protein 1 | Protein 2 | 10s | | 65s | | 129s | | | |
| | | *p*-value | Statistics | *p*-value | Statistics | *p*-value | Statistics | | |
| RS2Z32 (-LMB) | RS2Z32 (+LMB) | <0.0001 | **** | <0.0001 | **** | <0.0001 | **** | 32.8264% | 70.4650% |
| RS2Z32mutRNP1 (-LMB) | RS2Z32mutRNP1 (+LMB) | <0.0001 | **** | <0.0001 | **** | <0.0001 | **** | 40.3231% | 74.1742% |
| RS2Z32mutRNP2 (-LMB) | RS2Z32mutRNP2 (+LMB) | 0.4643 | ns | 0.9166 | ns | >0.9999 | ns | 42.3579% | 43.0882% |
| RS2Z32mutRRM (-LMB) | RS2Z32mutRRM (+LMB) | 0.0232 | * | 0.0002 | *** | <0.0001 | **** | 39.1768% | 64.4226% |
| RS2Z32mutZnK1 (-LMB) | RS2Z32mutZnK1 (+LMB) | 0.1261 | ns | 0.0089 | ** | 0.0016 | ** | 35.0049% | 52.6417% |
| RS2Z32mutZnK2 (-LMB) | RS2Z32mutZnK2 (+LMB) | 0.0021 | * | <0.0001 | **** | <0.0001 | **** | 39.4921% | 69.2022% |
| RS2Z32mutZnKs (-LMB) | RS2Z32mutZnKs (+LMB) | 0.0326 | * | <0.0001 | **** | 0.0001 | *** | 37.5356% | 62.0930% |
| RS2Z32mutRRM+ZnKs (-LMB) | RS2Z32mutRRM+ZnKs (+LMB) | 0.0758 | ns | 0.0016 | * | <0.0001 | **** | 34.7432% | 53.7743% |
| RS2Z33 (-LMB) | RS2Z33 (+LMB) | <0.0001 | **** | <0.0001 | **** | <0.0001 | **** | 30.8898% | 70.6307% |
| RS2Z33mutRNP1 (-LMB) | RS2Z33mutRNP1 (+LMB) | <0.0001 | **** | <0.0001 | **** | <0.0001 | **** | 34.2854% | 74.8619% |
| RS2Z33mutRNP2 (-LMB) | RS2Z33mutRNP2 (+LMB) | 0.6316 | ns | 0.9880 | ns | 0.8731 | ns | 41.7637% | 41.8250% |
| RS2Z33mutRRM (-LMB) | RS2Z33mutRRM (+LMB) | 0.0040 | ** | <0.0001 | **** | <0.0001 | **** | 39.6313% | 65.9904% |
| RS2Z33mutZnK1 (-LMB) | RS2Z33mutZnK1 (+LMB) | 0.0155 | * | <0.0001 | **** | 0.0002 | *** | 31.9430% | 48.3267% |
| RS2Z33mutZnK2 (-LMB) | RS2Z33mutZnK2 (+LMB) | <0.0001 | **** | <0.0001 | **** | <0.0001 | **** | 38.7004% | 68.7850% |
| RS2Z33mutZnKs (-LMB) | RS2Z33mutZnKs (+LMB) | 0.0002 | *** | <0.0001 | **** | <0.0001 | **** | 38.2748% | 59.3692% |
| RS2Z33mutRRM+ZnKs (-LMB) | RS2Z33mutRRM+ZnKs (+LMB) | 0.4194 | ns | 0.0495 | * | 0.0111 | * | 31.6903% | 43.2782% |
| RS2Z32 (-LMB) | RS2Z32mutRNP1 (-LMB) | 0.4856 | ns | 0.1024 | ns | 0.5946 | ns | 32.8264% | 40.3231% |
| RS2Z32 (-LMB) | RS2Z32mutRNP2 (-LMB) | 0.0377 | * | 0.0311 | * | 0.0163 | * | 32.8264% | 42.3579% |
| RS2Z32 (-LMB) | RS2Z32mutRRM (-LMB) | 0.6193 | ns | 0.1263 | ns | 0.3372 | ns | 32.8264% | 39.1768% |
| RS2Z32 (-LMB) | RS2Z32mutZnK1 (-LMB) | 0.9491 | ns | 0.5571 | ns | 0.8100 | ns | 32.8264% | 35.0049% |

| | | | | | | | | | |
|---|---|---|---|---|---|---|---|---|---|
| RS2Z32 (-LMB) | RS2Z32mutZnK2 (-LMB) | 0.3104 | ns | 0.1180 | ns | 0.0939 | ns | 32.8264% | 39.4921% |
| RS2Z32 (-LMB) | RS2Z32mutZnKs (-LMB) | 0.4467 | ns | 0.2553 | ns | 0.4102 | ns | 32.8264% | 37.5356% |
| RS2Z32 (-LMB) | RS2Z32mutRRM+ZnKs (-LMB) | 0.1683 | ns | 0.7511 | ns | 0.6299 | ns | 32.8264% | 34.7432% |
| RS2Z33 (-LMB) | RS2Z33mutRNP1 (-LMB) | 0.8024 | ns | 0.3975 | ns | 0.1480 | ns | 30.8898% | 34.2854% |
| RS2Z33 (-LMB) | RS2Z33mutRNP2 (-LMB) | 0.0048 | ** | 0.0043 | ** | 0.0066 | ** | 30.8898% | 41.7637% |
| RS2Z33 (-LMB) | RS2Z33mutRRM (-LMB) | 0.0204 | * | 0.0399 | * | 0.0174 | * | 30.8898% | 39.6313% |
| RS2Z33 (-LMB) | RS2Z33mutZnK1 (-LMB) | 0.2893 | ns | 0.7990 | ns | 0.6627 | ns | 30.8898% | 31.9430% |
| RS2Z33 (-LMB) | RS2Z33mutZnK2 (-LMB) | 0.0129 | * | 0.0386 | * | 0.0226 | * | 30.8898% | 38.7004% |
| RS2Z33 (-LMB) | RS2Z33mutZnKs (-LMB) | 0.0645 | ns | 0.0785 | ns | 0.0917 | ns | 30.8898% | 38.2748% |
| RS2Z33 (-LMB) | RS2Z33mutRRM+ZnKs (-LMB) | 0.2023 | ns | 0.8518 | ns | 0.2770 | ns | 30.8898% | 31.6903% |
| RS2Z32 (+LMB) | RS2Z32mutRNP1 (+LMB) | 0.0276 | * | 0.0878 | ns | 0.1077 | ns | 70.4650% | 74.1742% |
| RS2Z32 (+LMB) | RS2Z32mutRNP2 (+LMB) | 0.0019 | ** | <0.0001 | **** | <0.0001 | **** | 70.4650% | 43.0882% |
| RS2Z32 (+LMB) | RS2Z32mutRRM (+LMB) | 0.5485 | ns | 0.2139 | ns | 0.6284 | ns | 70.4650% | 64.4226% |
| RS2Z32 (+LMB) | RS2Z32mutZnK1 (+LMB) | 0.4819 | ns | 0.0024 | ** | 0.0030 | ** | 70.4650% | 52.6417% |
| RS2Z32 (+LMB) | RS2Z32mutZnK2 (+LMB) | 0.4479 | ns | 0.1577 | ns | 0.1633 | ns | 70.4650% | 69.2022% |
| RS2Z32 (+LMB) | RS2Z32mutZnKs (+LMB) | 0.7817 | ns | 0.0338 | * | 0.0455 | * | 70.4650% | 62.0930% |
| RS2Z32 (+LMB) | RS2Z32mutRRM+ZnKs (+LMB) | <0.0001 | **** | <0.0001 | **** | 0.0068 | ** | 70.4650% | 53.7743% |
| RS2Z33 (+LMB) | RS2Z33mutRNP1 (+LMB) | 0.3364 | ns | 0.1507 | ns | <0.0001 | **** | 70.6307% | 74.8619% |
| RS2Z33 (+LMB) | RS2Z33mutRNP2 (+LMB) | <0.0001 | **** | <0.0001 | **** | <0.0001 | **** | 70.6307% | 41.8250% |
| RS2Z33 (+LMB) | RS2Z33mutRRM (+LMB) | 0.1107 | ns | 0.2213 | ns | 0.3562 | ns | 70.6307% | 65.9904% |
| RS2Z33 (+LMB) | RS2Z33mutZnK1 (+LMB) | 0.0001 | *** | <0.0001 | **** | <0.0001 | **** | 70.6307% | 48.3267% |
| RS2Z33 (+LMB) | RS2Z33mutZnK2 (+LMB) | 0.1831 | ns | 0.6271 | ns | 0.5052 | ns | 70.6307% | 68.7850% |
| RS2Z33 (+LMB) | RS2Z33mutZnKs (+LMB) | 0.4542 | ns | 0.0011 | ** | <0.0001 | **** | 70.6307% | 59.3692% |
| RS2Z33 (+LMB) | RS2Z33mutRRM+ZnKs (+LMB) | <0.0001 | **** | <0.0001 | **** | <0.0001 | **** | 70.6307% | 43.2782% |

**Supplementary Table S8:** Statistical analyses of FLIP-shuttling assays of native proteins or mutant variants in the absence (−LMB) and upon leptomycin B (+LMB) treatment. Cells colored in light blue, light pink or light green represent, respectively, effect of leptomycin B treatment on the nucleocytoplasmic shuttling activity; effect of mutations on the nucleocytoplasmic shuttling activity, in absence of leptomycin B; and effect of mutations on the nucleocytoplasmic shuttling activity, in presence of leptomycin B (+LMB). For statistical analysis of normality, the D'Agostino and Pearson test was used. To calculate the significance of the differences between fluorescence intensities, an unpaired t test (parametric data) was performed. When at least one of the series of data failed the normality test, the comparison between the experiments were performed with the Wilcoxon signed-rank test. The differences observed were considered to be statistically significant if *p*-values were at least $< 0.05$, as indicated by asterisks (*P < 0.05, **P < 0.01, ***P < 0.001, ****P < 0.0001). n.s., not significant.

| qRT-PCR | | | | |
|---|---|---|---|---|
| **Gene ID** | **Symbol** | **Forward (5' – 3')** | **Reverse (5' – 3')** | **Reference** |
| AT3G53500 | RS2Z32 | CAAATCGCTACCGTTGAATCTC | AAGGAAGTGAACCGCGGAT | This study |
| AT2G37340 | RS2Z33 | AACAGCCCCAAGAAGCTTAGG | AGCTTCGGCTACGGCTAAGACT | This study |
| AT1G58050 | AT1G58050 | CCATTCTACTTTTTGGCGGCT | TCAATGGTAACTGATCCACTCTGATG | (Rausin *et al.*, 2010) |

**Supplementary Table S9:** Primers used for mRNA levels analysis (qRT-PCR).

| Native RRM of RS2Z32 | | | | |
|---|---|---|---|---|
| **#** | **Sequence** | **Length (bp)** | **Purine (%)** | **Pyrimidine (%)** |
| **1** | UAAGACCGGUAUCCCAACCCCGUA | 24 | 45.83 | 54.17 |
| **2** | UAAACAGACCGGUAUCCCUUAGCA | 24 | 50.00 | 50.00 |
| **3** | UAAGUCCGGUACCCCCUAAA | 21 | 42.86 | 57.14 |
| **4** | ACCAGCAUCGAUAGAGCUACUCCA | 24 | 50.00 | 50.00 |
| **5** | CCGAGCGGUAUCCCACAUGAU | 21 | 47.62 | 52.38 |
| **6** | UCCCCUCGUCUCUAUAUAUUCCGCUAUUCUCAUAUGCC | 38 | 23.68 | 76.32 |
| **7** | GGGCGGGUUUAGGCCGUUGGGAAC | 24 | 62.50 | 37.50 |
| **8** | GAACUUAUGAAGAGUAAAAAUAUCGAGACGAGGGG | 35 | 74.29 | 25.71 |
| **9** | GAUGAUCAUCCGUGGAAUUGGAA | 23 | 60.87 | 39.13 |
| **10** | AUAUCGACAAAUAGGGAG | 18 | 72.22 | 27.78 |

**Supplementary Table S10:** List of sequences submitted to MEME to find a significant consensus for the native RRM of RS2Z32.

| Native RRM of RS2Z33 | | | | |
|---|---|---|---|---|
| **#** | **Sequence** | **Length (bp)** | **Purine (%)** | **Pyrimidine (%)** |
| **1** | AAGUAGCUUCCGGUAUCCCUUA | 22 | 40.91 | 59.09 |
| **2** | UGUAGAGCCGGUAUCCCAGCUGUC | 24 | 45.83 | 54.17 |
| **3** | AGAGCAAUCUGCUAACUCCCUGG | 23 | 47.83 | 52.17 |
| **4** | UACAAUCUGCUACUCCCUGUAC | 22 | 31.82 | 68.18 |
| **5** | UUACAACCGGUAUCCCAGCUACCG | 24 | 41.67 | 58.33 |
| **6** | CUGAUCCGGUAUCCUAAACCGACA | 24 | 45.83 | 54.17 |
| **7** | AUGAUCCGGUAUCCUAACCCGACA | 24 | 45.83 | 54.17 |
| **8** | GCGACGGGGGACAAGAGGGUGUGA | 24 | 79.17 | 20.83 |
| **9** | GCAUAAUAAAGGAAGGCGGGGAUA | 24 | 79.17 | 20.83 |
| **10** | GGAGAAAUUAAUACGACUCACUAUAGGGAG | 30 | 66.67 | 33.33 |

**Supplementary Table S11:** List of sequences submitted to MEME to find a significant consensus for the native RRM of RS2Z33.

| Mutated RRM of RS2Z32 | | | | |
|---|---|---|---|---|
| **#** | **1st set of sequences (Motif #1)** | **Length (bp)** | **Purine (%)** | **Pyrimidine (%)** |
| **1** | GCGGGCGGGUUAUCA | 15 | 60.00 | 40.00 |
| **2** | AAAGAGGGGUGC | 12 | 83.33 | 16.67 |
| **3** | AUAUCGACAAAUAGGGAG | 18 | 72.22 | 27.78 |
| **4** | GACGAUGCGCAUAAACAAGAUGGUG | 25 | 68.00 | 32.00 |
| **5** | GAAAUGUGGGCUCAGGGAGUUGGUA | 25 | 68.00 | 32.00 |
| **6** | GCCACAGUAUUUGAGAGA | 18 | 61.11 | 38.89 |
| **7** | AACUGACAACGCAUCUUAACGAAUA | 25 | 56.00 | 44.00 |
| **8** | UAAGUCCGGUACCCCCCUAAA | 21 | 42.86 | 57.14 |
| **9** | GGAAAAUUAAUACGACUCACUAUAGGGAG | 29 | 65.52 | 34.48 |
| **10** | GCGGGCUAUAAAGUAUGAAGCGGCU | 25 | 64.00 | 36.00 |

| Mutated RRM of RS2Z32 | | | | |
|---|---|---|---|---|
| **#** | **2nd set of sequences (Motif #2)** | **Length (bp)** | **Purine (%)** | **Pyrimidine (%)** |
| **1** | CGGGAAAGAGGGUUAC | 16 | 75.00 | 25.00 |
| **2** | CAAACGGGUACUACAUAUUGUGUA | 24 | 54.17 | 45.83 |
| **3** | CGGGGGUAGCGGGA | 14 | 78.57 | 21.43 |
| **4** | GAGGGGAACCGGGACGGGAGUGAA | 24 | 83.33 | 16.67 |
| **5** | GACAGAAGACGAGGGAG | 17 | 88.24 | 11.76 |
| **6** | GACAGACGACGAG | 13 | 76.92 | 23.08 |
| **7** | GAAAGGUUAAGGGGGGGUUUGAUC | 24 | 70.83 | 29.17 |
| **8** | AUAGUGAGUCGUAAUAAUAUCCGACUCACUAUA | 33 | 51.52 | 48.48 |
| **9** | GGAGGGAGGACGAUGCGGGG | 20 | 85.00 | 15.00 |
| **10** | GUUAAAGCAUGGUGCCACUGCGAA | 24 | 58.33 | 41.67 |

**Supplementary Table S12:** List of sequences (two sets of ten sequences) submitted to MEME for the mutated RRM of RS2Z32.

| Mutated RRM of RS2Z33 | | | | |
|---|---|---|---|---|
| **#** | **1st set of sequences (Motif #1)** | **Length (bp)** | **Purine (%)** | **Pyrimidine (%)** |
| **1** | GACGGGCCUUAGUGAGUGAUGCU | 23 | 56.52 | 43.48 |
| **2** | GCGGGCCAUGAUACAAAGCGCGCU | 24 | 58.33 | 41.67 |
| **3** | CGGGGAAUUGAGCGAGGGGCAGA | 23 | 78.26 | 21.74 |
| **4** | AAAAAACAAGGGCGUAAGGCGGG | 23 | 82.61 | 17.39 |
| **5** | UACAAGACACCAUUUGUGGCCCGC | 24 | 45.83 | 54.17 |
| **6** | AGGGGGGGAGGAUAUCAAAUGGG | 23 | 82.61 | 17.39 |
| **7** | UCCUGACACACUAUGGGAGGG | 21 | 57.14 | 42.86 |
| **8** | AGGACGAUGCGGUCCUGA | 18 | 61.11 | 38.89 |
| **9** | GGUUAUCGAAAACACUAGGGGGGG | 24 | 70.83 | 29.17 |
| **10** | UAUAAUUUAAGCUACCUGUAGCUA | 24 | 45.83 | 54.17 |

| Mutated RRM of RS2Z33 | | | | |
|---|---|---|---|---|
| **#** | **2nd set of sequences (Motif #2)** | **Length (bp)** | **Purine (%)** | **Pyrimidine (%)** |
| **1** | GCCGGGCAGGGGAACGGGUGAAUA | 24 | 75.00 | 25.00 |
| **2** | CUCACUAUAGGGAGGACGAUGCGAGC | 26 | 61.54 | 38.46 |
| **3** | AAAAACCGAAACCAACAAAAAAAAC | 25 | 76.00 | 24.00 |
| **4** | AGCAGAUACCGACGGAGGACAAGGCGGGG | 29 | 75.86 | 24.14 |
| **5** | GGAAGAUUUUGGAAGAAGGGGUCA | 24 | 75.00 | 25.00 |
| **6** | GCGGGAAAAUUGAGGAAAUACGUAC | 25 | 72.00 | 28.00 |
| **7** | GGGAAAGAAUCAGGCGGGAAUUAA | 24 | 79.17 | 20.83 |
| **8** | AUUAUACCUCCAGGGUUAUAGAAUU | 25 | 48.00 | 52.00 |
| **9** | ACGAAGGGGAGGACGAUGAA | 20 | 85.00 | 15.00 |
| **10** | CAAACAAGGGUGGGUAAACGGAGA | 24 | 79.17 | 20.83 |

**Supplementary Table S13:** List of sequences (two sets of ten sequences) submitted to MEME for the mutated RRM of RS2Z33.

| Native RRM+ZnKs of RS2Z32 | | | | |
|---|---|---|---|---|
| **#** | **Sequence** | **Length (bp)** | **Purine (%)** | **Pyrimidine (%)** |
| **1** | CAAAAAAUCACAGGAGGUGAAGGA | 24 | 79.17 | 20.83 |
| **2** | GGUAAGGGUGUGAAAUAAGUGCUC | 24 | 66.67 | 33.33 |
| **3** | GCGAUGAUGAUGUAUCGCCAAACU | 24 | 54.17 | 45.83 |
| **4** | UUAAAAGAGGUGGCAUGUUUAAGA | 24 | 66.67 | 33.33 |
| **5** | CUUACGAGAGUGGUGAACCUGGCU | 24 | 54.17 | 45.83 |
| **6** | GCAGUGGGCUAAGGGAG | 17 | 76.47 | 23.53 |
| **7** | AGGAGCCACAUCAGAAUAAGCG | 22 | 68.18 | 31.82 |
| **8** | UGUGGAGGAUGUGCAGUUAGCCCG | 24 | 58.33 | 41.67 |
| **9** | GGUGGGAUUGAGUCACGUUUAACA | 24 | 58.33 | 41.67 |
| **10** | GACUGAAUGUUGGCAUGUUAG | 21 | 57.14 | 42.86 |
| **11** | GCAUACACAAAGGUGAAGUGUAACUAGACG | 30 | 66.67 | 33.33 |
| **12** | GAAUUGAUAUCAGAGGGAGGCGAUGCGGGAAUUGAUAU | 38 | 68.42 | 31.58 |
| **13** | UGAGGAGGGUUUGACAGCAGUA | 22 | 68.18 | 31.82 |
| **14** | GUGGGACUAUGAAAGAUAUACGGC | 24 | 66.67 | 33.33 |
| **15** | UAGAAGUGAAGGUUGAUAAAGGU | 23 | 73.91 | 26.09 |
| **16** | CGGGAGGGAUACGGAGAUAUCUCG | 24 | 66.67 | 33.33 |

| | | | | |
|---|---|---|---|---|
| **17** | GCCAGGGAGGAGUAGAUGUAGUGA | 24 | 75.00 | 25.00 |
| **18** | AAGAUAUGGGAGGGCAGGAUUAGA | 24 | 79.17 | 20.83 |
| **19** | GGGAGCAUGGGGAAAUGGGCGCU | 23 | 73.91 | 26.09 |
| **20** | CUCCCUAUAGUGAGUCGUAUUAAUUUCG | 28 | 39.29 | 60.71 |
| **21** | AAAACAAAGUGUGGAAUGAUGGCU | 24 | 70.83 | 29.17 |
| **22** | UUGUGGCAUGUCCUCCGGCUUUAA | 24 | 37.50 | 62.50 |
| **23** | AUCUGGGCGGGGUUAUGGGUAUAA | 24 | 62.50 | 37.50 |
| **24** | UAAAAUGGUGAGCCCGCAAUGCAU | 24 | 58.33 | 41.67 |
| **25** | GGGGGUUCGUAGCAGGGGGCUAAU | 24 | 66.67 | 33.33 |
| **26** | UGGGUAAGACGGAUACGGGAGGUU | 24 | 70.83 | 29.17 |
| **27** | AUAAUGUACGUUAGAAGUAAAGGU | 24 | 66.67 | 33.33 |
| **28** | CGGGAGGGUGAAAGGAUGGGGUAA | 24 | 83.33 | 16.67 |
| **29** | CCGAUGUAUGAACAGAUCCGAUUG | 24 | 54.17 | 45.83 |
| **30** | UGCCUACCCAUGGGAGGACUCG | 22 | 50.00 | 50.00 |

**Supplementary Table S14:** List of sequences submitted to MEME to find a significant consensus for the native RRM+ZnKs of RS2Z32.

| Native RRM+ZnKs of RS2Z33 | | | | |
|---|---|---|---|---|
| **#** | **Sequence** | **Length (bp)** | **Purine (%)** | **Pyrimidine (%)** |
| **1** | GACAUCAACAAAGGGUCAUAUUGA | 24 | 62.50 | 37.50 |
| **2** | GAUGCUCAUAACACAAGGGGGAAC | 24 | 66.67 | 33.33 |
| **3** | GAGGGAAUGUAGAUGAGUAGGGG | 23 | 82.61 | 17.39 |
| **4** | AAGGCGAGGAGAAGUAUGUGCAACU | 25 | 72.00 | 28.00 |
| **5** | GCGGGAGAGGGAGGACGAUG | 20 | 85.00 | 15.00 |
| **6** | GCGGGAGAUAUAAGAGCGUUUGG | 23 | 69.57 | 30.43 |
| **7** | UCAAGUAGCAUCACCAAUGAAAUU | 24 | 54.17 | 45.83 |
| **8** | GGUGUGGGAAGAAAUGAGAGG | 21 | 85.71 | 14.29 |
| **9** | AUGACACAAAAUAUAAUGUGAUUU | 24 | 58.33 | 41.67 |
| **10** | AGUUAACAUAACCAGGGAUGCAAU | 24 | 62.50 | 37.50 |
| **11** | AGAUUUAGAGAGGAUUACCCA | 21 | 61.90 | 38.10 |
| **12** | UAUUAAAGGCUGCCUCCACCACCA | 24 | 41.67 | 58.33 |
| **13** | GAAGAGGGAAGACAGGAGGGUAAG | 24 | 91.67 | 8.33 |
| **14** | AUCAUCAAUACGAUAAUAUUAUG | 23 | 52.17 | 47.83 |
| **15** | ACUACGGUUCCGCAGAGGGGAAU | 23 | 60.87 | 39.13 |
| **16** | GGGAAUCCGAACCUAAGAGGAGC | 23 | 69.57 | 30.43 |

| | | | | |
|---|---|---|---|---|
| **17** | UUAGACGAGAGCGGGA | 16 | 75.00 | 25.00 |
| **18** | GCCAACGGGCCAAUUGAGAGACCG | 24 | 62.50 | 37.50 |
| **19** | GGAAAGAGCGAAACAGGGCGCUCA | 24 | 75.00 | 25.00 |
| **20** | UAACCGCUACAUGCACAGACGGGA | 24 | 58.33 | 41.67 |
| **21** | GAGGAACCUGACGCAGACAGAACG | 24 | 70.83 | 29.17 |
| **22** | ACGAAUGAAAGGGUCUAGAUUCAG | 24 | 66.67 | 33.33 |
| **23** | AUAGCGACUGGCGGAGGGUGUCAA | 24 | 66.67 | 33.33 |
| **24** | ACGGAGAAUGCUAACUUUACCCCG | 24 | 50.00 | 50.00 |
| **25** | CUUUAAUAGCAACCUCCGAUAGUA | 24 | 45.83 | 54.17 |
| **26** | AAAAGAGGGGUAACCAGUGAGGG | 23 | 82.61 | 17.39 |
| **27** | UUAGACGACGGGGA | 14 | 71.43 | 28.57 |
| **28** | ACAGACGACGAGCGGGACAGACG | 23 | 73.91 | 26.09 |
| **29** | GUGGCAUGACGUUAGGGGAGGAUU | 24 | 66.67 | 33.33 |
| **30** | CUAGGCAAGCCAUCAGUUAGAGCA | 24 | 58.33 | 41.67 |

**Supplementary Table S15:** List of sequences submitted to MEME to find a significant consensus for the native RRM+ZnKs of RS2Z33.

Califice, S., D. Baurain, M. Hanikenne and P. Motte (2012). A single ancient origin for prototypical serine/arginine-rich splicing factors. *Plant Physiology* **158**(2): 546-560.

De Franco, S., J. Vandenameele, A. Brans, O. Verlaine, K. Bendak, C. Damblon, A. Matagne, D. J. Segal, M. Galleni, J. P. Mackay and M. Vandevenne (2019). Exploring the suitability of RanBP2-type Zinc Fingers for RNA-binding protein design. *Scientific Reports* **9**(1): 2484.

Fanara, S., M. Schloesser, M. Joris, S. De Franco, M. Vandevenne, F. Kerff, M. Hanikenne and P. Motte (2024). The Arabidopsis SR45 splicing factor bridges the splicing mc fachinery and the exon-exon junction complex. *Journal of Experimental Botany* **75**(8): 2280-2298.

Maris, C., C. Dominguez and F. H. Allain (2005). The RNA recognition motif, a plastic RNA-binding platform to regulate post-transcriptional gene expression. *The FEBS Journal* **272**(9): 2118-2131.

Rausin, G., V. Tillemans, N. Stankovic, M. Hanikenne and P. Motte (2010). Dynamic nucleocytoplasmic shuttling of an Arabidopsis SR splicing factor: role of the RNA-binding domains. *Plant Physiology* **153**(1): 273-284.

Stankovic, N., M. Schloesser, M. Joris, E. Sauvage, M. Hanikenne and P. Motte (2016). Dynamic Distribution and Interaction of the Arabidopsis SRSF1 Subfamily Splicing Factors. *Plant Physiology* **170**(2): 1000-1013.
